# Female oviposition decisions are influenced by the microbial environment

**DOI:** 10.1101/2024.07.03.601843

**Authors:** Emily K. Fowler, Lucy A. Friend, Emily R. Churchill, Douglas W. Yu, Marco Archetti, Andrew F.G. Bourke, Amanda Bretman, Tracey Chapman

## Abstract

In ovipositing animals, egg placement decisions can be key determinants of offspring survival. One oviposition strategy reported across taxa is egg clustering, whereby a female lays multiple eggs next to one another or next to the eggs of other females. The fitness benefits of egg clustering, especially in mixed maternity clusters, are unknown. In some species, mothers provision eggs with diffusible defence compounds, such as antimicrobials, raising the possibility of public good benefits arising from egg clustering. Here we report that *Drosophila melanogaster* females frequently lay eggs in mixed maternity clusters. We tested two hypotheses for potential drivers of this oviposition behaviour: (i) the microbial environment affects fecundity and egg placement in groups of *D. melanogaster* females; (ii) *D. melanogaster* eggs exhibit antimicrobial activity. The results partially supported the first hypothesis. Females exposed to environmental microbes that naturally colonised the oviposition substrates in the absence of antimicrobial preservatives reduced their levels of fecundity but did not significantly alter egg clustering. In contrast, the presence of commensal (fly-associated) microbes did not affect oviposition. The second hypothesis was not supported. There was no evidence of antimicrobial activity, either in whole eggs or in soluble surface material extracted from them. In conclusion, while there was no evidence that oviposition decisions are shaped by the opportunity to share antimicrobials, there is evidence that the microbial environment provides cues that females use to make sophisticated decisions on fecundity and egg placement.

## Introduction

For an oviparous animal, deciding where and how to place eggs can have major fitness consequences for parents and their offspring. Both abiotic and biotic factors at the site of oviposition can determine the survival, development and phenotype of offspring (1). Consistent with this, oviposition site choice is non-random across many taxa. Several explanatory hypotheses have been proposed, (reviewed in ref. 2). Of these, maximising embryo survival has been viewed as one of the most important factors, whereby females are expected to choose sites that minimize predation and competition for resources and optimise abiotic conditions for embryo development and hatching. For example, the tree-hole breeding frog *Phrynobatrachus guineensis* oviposits in temporary pools and exhibits a preference for sites with the appropriate level of water persistence required for successful offspring development. This species also prefers to oviposit at sites already inhabited by conspecific eggs and tadpoles, despite a negative relationship between total tadpole density and size at metamorphosis, and it is thought the presence of conspecifics may indicate, or result in, lower predation risk (3). Similarly, in the pine sawfly *Neodiprion sertifer*, female sawflies prefer to oviposit on trees with high resin acid concentrations, which results in offspring reaching a smaller pupal weight but lowers the vulnerability to attack by parasitoids (4).

Furthermore, animals can exhibit oviposition decisions within a single oviposition patch or substrate by adjusting the number of eggs they lay through delaying oviposition if the substrate or environmental conditions are perceived as sub-optimal. Individuals can also position their eggs in non-random patterns. For example, females can lay their eggs singly or cluster their eggs together, sometimes with eggs of other females. Egg clustering behaviour, including mixed-maternity clustering, has been reported for many taxa including reptiles and amphibians (5), birds (6), fish (7) as well as in several invertebrate species (e.g. Refs. 8, 9). Egg clustering also occurs in the fruit fly *Drosophila melanogaster* and, furthermore, egg clustering is a plastic behaviour which increases in frequency with social density (10).

Several fitness benefits of egg clustering have been proposed, although empirical evidence to support them remains scant (11). For example, clustering eggs could be the outcome of females reducing site and substrate evaluation times and instead relying on the decisions of others (9). Alternatively, clustering could reduce egg predation risk if predators have limits on their search or consumption time or capacity. For instance, *Iphiseius degenerans* mites oviposit in clusters in the tufts of leaf hairs (acarodomatia). Female mites prefer to cluster their eggs in acarodomatia already containing eggs, and clustered eggs were less likely to be predated by thrips, relative to when eggs are scattered across acarodomatia (8). Egg clustering may also increase egg survival during exposure to abiotic factors, such as low humidity. For example, in the Nymphalid butterfly *Chlosyne lacinia*, hatching success is positively related to humidity, and eggs clustered in larger groups have greater desiccation resistance in comparison to small groups of monolayered eggs (12).

In this study, we propose an additional hypothesis - that clustered eggs benefit from the collective increased concentrations of defensive (i.e. antimicrobial) compounds potentially provisioned to the egg surface by the mother (13). When the defensive compounds are external and diffusible, they are potential ‘public goods’, such that eggs without the compounds nonetheless receive benefits from the compounds released by nearby eggs (14). For example, Mediterranean fruitfly (*Ceratitis capitata*) females smear the surface of their eggs with secretions containing ceratotoxins - a family of broad-acting antimicrobial peptides (AMPs), which are produced in the female reproductive tract (15). Though the genes encoding ceratotoxins have no known homologues outside *Ceratitis* (16), some AMPs of *Drosophila* are similarly expressed in the female reproductive tract and thus also have the potential to be transferred extracellularly to the egg surface. For example, the anti-fungal peptide encoding gene *Drosomycin* is expressed in the reproductive tract epithelium (17, 18) and the anti-bacterial encoding gene *Drosocin* is constitutively expressed in the female oviduct (18, 19). The promoters of other AMP genes including *cecropin*, *defensin*, *metchnikowin* and *attacin* are also active in the reproductive tract (18). Furthermore, a transcriptomic study of female reproductive tissues found that some AMP genes were upregulated following mating (20). It is not yet known why this upregulation occurs, but one possibility is to enable the production of higher amounts of AMPs to protect the elevated numbers of eggs that are produced and laid following mating.

Consistent with the hypotheses we test in this study is the extensive evidence that the microbial environment influences oviposition behaviour of insects, including *Drosophila*. For example, when offered a direct choice between substrates containing commensal microbes (i.e. members of the fly-associated microbiome) vs. sterile substrates, *D. melanogaster* prefer to lay on microbe-inoculated substrates, whereas *D. suzukii* prefers sterile substrates (21). These differences may reflect the natural oviposition substrates of these two species, with *D. melanogaster* laying into fermenting fruit and *D*. *suzukii* into ripening fruit. The Oriental fruitfly *Bactrocera dorsalis* uses a volatile compound associated with the presence of egg-surface bacteria to avoid laying into fruits already occupied by conspecific eggs (22). There is also evidence that *D. melanogaster* uses sucrose levels as a means of assessing the presence or level of commensal bacteria in their food, since the lactic acid bacteria *Enterococci* metabolises and therefore depletes sucrose within food sources (23). *D. melanogaster* eggs also appear to be dependent on microbes for successful development, with germ-free eggs failing to develop beyond the second instar larvae when reared in food lacking yeast (23, 24).

*D. melanogaster* females lay eggs in decomposing fruit with a rich microbial environment that is very likely to contain a mix of beneficial, neutral and pathogenic microbial species (25, 26). Although few extracellular pathogens have so far been identified as attacking *D. melanogaster* eggs (26), ingestion of some bacterial strains by larvae can be fatal (25). This suggests there should be selection for choosing or maintaining pathogen-free oviposition sites. Consistent with this, *D. melanogaster* females can detect and avoid the odorous compound geosmin, which is produced by some microbes, including pathogenic species (26). Collectively, these data support the hypothesis that female flies choose oviposition sites according to the prevailing microbial milieu and/or protect their eggs from pathogens by deploying antimicrobials. The latter raises the possibility that oviposition clustering decisions are shaped by potential public good benefits for antimicrobial protection. The aims of this study were to investigate these ideas by testing the hypotheses that: (1) *D. melanogaster* females plastically adjust egg placement based on the microbial environment; and (2) *D. melanogaster* eggs exhibit broad spectrum antimicrobial activity.

## Materials and methods

### Fly stocks and handling

Wild type *D. melanogaster* flies were sourced from a large laboratory population originally collected in the 1970s in Dahomey (Benin) and maintained in stock cages with overlapping generations. Flies carrying the *scarlet* mutation were maintained in the same way. Flies were reared on standard sugar yeast agar (SYA) medium (100 g brewer’s yeast (*MP Biomedicals*, *Fisher Scientific #11425722*), 50 g white caster sugar (*Tate & Lyle*), 15 g agar (*Formedium #AGA01*), 30 ml Nipagin (methylparaben, 10% w/v solution, dissolved in 95% Ethanol), and 3 ml propionic acid (*Sigma-Aldrich #P5561*), per litre of medium) in a controlled environment (25°C, 50% humidity, 12:12 hour light:dark cycle). Eggs were collected from population cages on grape juice agar plates (50 g agar, 600 ml red grape juice (medium dry red wine kit, *Magnum*), 42 ml 10% w/v Nipagin solution per 1.1 l RO H_2_O) supplemented with fresh yeast paste (Saf-levure active dry yeast, *Lesaffre*), and first instar larvae were transferred to SYA medium at a standard density of 100 per vial (glass, 75 × 25 mm, each containing 7 ml medium). Male and female adults were separated within 6 hours of eclosion under ice anaesthesia and stored in single sex groups of 10/vial.

### Statistical methods

All statistical analyses were conducted using R version 4.2.1 (27). Graphs were produced using *ggplot2* (28) and *ggpubr* (29) packages. Summary statistics were produced using the *Rmisc* package (30). We defined an egg cluster as a group of two or more eggs where any part of the main body of an egg was in physical contact with any part of the main body of another egg (Figure S1). Egg clustering proportion was calculated for vials containing ≥ 2 eggs. Clustering proportion per vial was calculated as the number of all eggs in any cluster divided by the total number of eggs (Figure S1). For all analyses, full models containing all explanatory variables and their interactions were fitted in the first instance. Non-significant interactions (as tested using the anova function) were then removed from the models using a stepwise process. Model residuals were plotted and checked visually, using the DHARMa package where possible (31). Overdispersed or zero-inflated models were refitted using quasi-, negative binomial or hurdle GLMs as described below. Final model outputs are presented in the supplementary material. Details of specific analyses are given in each section below.

### Hypothesis 1

#### Effect of environmental microbes and nutrients on oviposition

To test the effect of environmental microbes (i.e. microbes occurring naturally in the environment, which colonise substrates in the absence of sterilization or preservatives) and the nutritional content of the oviposition substrate on egg placement, we conducted oviposition assays on low or standard nutrient media in the presence or absence of antimicrobial preservatives. Unmated females were collected as described above and placed into groups of 4 in fresh SYA vials under CO_2_ anaesthesia at ∼4 days post-eclosion. After 2 days, groups of 6 males were introduced to female vials and left for 2 hours to mate before the females were transferred to new vials containing one of four substrates: 1) standard SYA; 2) standard SYA lacking the preservatives propionic acid and Nipagin (methyl paraben); 3) low nutrient SYA, with 25% of the yeast and 25% of the sugar of standard SYA; and 4) low nutrient SYA lacking the preservatives propionic acid and Nipagin. When preservatives were omitted, the equivalent volume of RO water was added instead. In total, 30 vials of females were set up for each treatment level. Females were allowed to oviposit for 3-4 hours before they were removed. The number of eggs laid, and the number and size of egg clusters (defined as ≥2 eggs in physical contact), were recorded immediately. The number of hatched eggs was recorded in each vial 48 hours later. The number of pupae in each vial was recorded 7 days following oviposition and the total number of adult offspring was recorded 11-12 days after oviposition.

To test the effect of substrate condition (low nutrient SYA ± preservatives, n = 30 per treatment) on the extent of mixed-maternity egg clustering (≥ 1 egg in direct contact with ≥ 1 egg laid by ≥ 2 different females), we followed the same protocol described above but used an oil-based dye to stain non-focal females and consequently their eggs. This allowed us to distinguish eggs laid by the focal female from those of three non-focal females. The non-focal females carried the *scarlet* mutation (a recessive eye colour marker) and were reared from first instar larvae on SYA containing 1400 ppm Sudan Black B dye (*Sigma-Aldrich* #199664) dissolved in corn oil (*Mazola*) but otherwise treated the same as focal females. The focal females were from the wild-type Dahomey stock and treated as described above. Non-focal females were mated to *scarlet* males prior to the oviposition assay. The *scarlet* phenotype allowed us to distinguish and score adult offspring of the focal and non-focal females.

We analysed the effect of nutrient level and preservative presence on the total number of eggs laid using a two-part hurdle model from the *pscl* package (32). The probability of eggs being laid was modelled with a binomial distribution and logit link function, while a zero truncated negative binomial distribution with log link function was used for the count part of the model. Nutrient (2 levels: low, standard) and preservative (2 levels: absent, present) were fixed factors in both parts of the model. We analysed the effect of nutrient level and preservative presence on egg clustering proportion, egg hatchability, egg to pupa viability, egg to adult viability and hatched egg to adult viability using quasibinomial GLMs. The variable “total eggs” was included as a fixed factor when modelling clustering proportion as a response variable, and “clustering proportion” was included as a fixed factor when modelling egg hatchability as a response. There was significant collinearity between “total eggs” and “clustering proportion” as measured using a Pearson’s correlation test from the *performance* package (33). Therefore, for all other measures of development, two separate models were run per response variable – one model included total eggs as a fixed factor and the other included clustering proportion. All models included nutrient and preservative as fixed factors. Reported significance values were derived using the Anova (Type II) function from the *car* package (34).

To test the reliability of egg maternity scoring, we used the one-way intraclass correlation coefficient from the *irr* package (35) to test the agreement between focal (non-dyed) eggs counted and the number of focal (wildtype) offspring which eclosed from each vial. We also generated a Bland Altman Plot of focal eggs and focal offspring using the *BlandAltmanLeh* package (36).

#### Effect of antimicrobial preservatives alone on oviposition

To test the effect of antimicrobial preservatives on oviposition in the absence of microbes, we provided females with sterile oviposition substrates that contained or lacked individual preservatives. This experiment differed from the first because all substrates were sterile, and thus females were not exposed to environmental microbes at the start of the oviposition assay, regardless of the presence of preservatives. Additionally, we tested the response of females to each individual antimicrobial preservative (Nipagin, ethanol or propionic acid). Unmated females were collected as described above and placed into groups of four on standard SYA food at ∼4 days old. After 2 days, groups of 6 males were introduced to female vials and left for ∼2 hours to mate. Females were then moved in their groups onto one of five oviposition substrates in glass vials (30 vials per treatment). All oviposition media was autoclaved for sterilisation. The test preservatives, or sterilised water, were added to the media after autoclaving. The different media were dispensed into sterile vials, under sterile conditions (inside an airflow cabinet) and topped with sterile cotton wool. The five test media (all sterile) were: (1) Standard – SYA including both Nipagin and propionic acid as per the standard 100% yeast and sugar recipe (described above); (2) No preservatives: both Nipagin and propionic acid omitted; (3) Propionic acid only – Nipagin omitted; (4) Nipagin only – propionic acid omitted; (5) Ethanol only – 95% ethanol added instead of Nipagin; propionic acid omitted. Where one or more preservatives were omitted, the equivalent volume of sterile RO water was added instead. Females were allowed to oviposit for 3-4 hours. The total numbers of eggs and egg clusters, and of resulting offspring, were scored for each vial as described above.

We analysed the effect of antimicrobial preservatives on total eggs using a negative binomial GLM, and we analysed the effect of preservatives and total eggs on egg clustering proportion using a quasibinomial GLM. Preservatives (5 levels: standard; no preservatives; propionic acid only; Nipagin only; ethanol only) was included as a fixed factor in all models. Again, because of collinearity between total eggs and clustering proportion, when analysing the effect of preservatives on egg to adult viability we ran two separate quasibinomial GLMs. One model included preservatives and total eggs as fixed factors and the other included preservative and clustering proportion as fixed factors. All reported significance values were derived using the Anova (Type II) function from the *car* package.

#### Effect of commensal and pathogenic microbes on oviposition

To test the effect of fly-associated microbial communities on egg placement in the absence of preservatives, we used oviposition substrates lacking preservatives, with or without microbial washes added to the surface of the substrate. Females were collected, stored and mated as described for the two experiments described above. After mating, groups of four females were moved onto one of four different oviposition substrates, with 30 vials per treatment. All oviposition media were autoclaved and dispensed under sterile conditions as described above, and all media lacked preservatives. Twenty-four hours before the oviposition assay, each oviposition substrate was spiked with 40 µl of one of four different washes - negative control, fly background control, commensal microbes (i.e. members of the fly-associated microbiome), and a culture of the bacteria *Alcaligenes faecalis* M3A. *Alcaligenes faecalis* is an identified pathogen of *Drosophila melanogaster* (*25*). The commensal microbe and fly background washes were made by placing 3 sterile grape juice agar plates into mini-cages with 300 adult flies per cage (1:1 sex ratio) for 10 hours. The flies were then discarded, and each plate was repeatedly washed with 2.5 ml sterile RO H_2_O. Half of this wash was used as the commensal microbe treatment, and the other half was filter sterilised to generate the fly background control (*Corning Costar* Spin-X centrifuge tube filter, 0.45 µm pore size, #8163). To generate the negative control wash, 2.5 ml RO H_2_O was used to wash the surface of 3 separate sterile grape juice agar plates that remained unexposed to flies, and the entirety of this wash was filter sterilised to remove any microbial contaminants. Finally, an overnight culture of the gram-negative bacterium *Alcaligenes faecalis* M3A was inoculated 1:100 into 100 ml Lysogeny Broth (see below for recipe) and grown at 30°C, 200 RPM for 3 hours, resulting in an optical density of 0.14 at 600 nm wavelength. 1 ml of this culture was centrifuged for 2 mins at 15,000 RPM and resuspended using 2 ml of the negative control wash to create the *A. faecalis* treatment. Following addition of the washes to the oviposition surfaces, vials were incubated for 24 hours at 25°C before the oviposition assay. Females were allowed 3-4 hours to lay eggs, as for the other oviposition assays. A set of 5-6 unexposed vials from each treatment was incubated at 25°C for the duration of the experiment. These vials were spiked with the washes, but never exposed to flies. This enabled us to check the extent of microbial growth from the washes, separate to the microbes introduced by females during the oviposition assay.

We analysed the effect of microbes on total eggs using a negative binomial GLM and analysed the effect of microbes and total eggs on egg clustering proportion using a quasibinomial GLM. Microbes (4 levels: negative control, fly background control, fly commensal microbes, *A. faecalis*) was included as a fixed factor in all models. As for the previous two experiments, we ran two separate quasibinomial GLMs for analysing effect of microbes on egg to adult viability. One model included microbes and total eggs as fixed factors, and the other included microbes and clustering proportion as fixed factors. All reported significance values were derived using the Anova (Type II) function from the *car* package.

### Hypothesis 2

#### Antimicrobial activity of egg surface molecules

To test if *D. melanogaster* eggs exhibit antimicrobial activity, we conducted radial diffusion assays (37) using whole eggs, or soluble material washed from the surface of laid eggs against the bacteria *Escherichia coli* dh5α, *Alcaligenes faecalis* M3A and *Micrococcus luteus* and the yeast *Saccharomyces cerevisiae* NYCC 505. As a positive control, we tested whole eggs and soluble material washed from the eggs of the Toliman strain of Mediterranean fruit fly (*Ceratitis capitata*), since Medfly eggs are known to exhibit antimicrobial activity (15). Wildtype *Ceratitis capitata* flies of the Toliman strain were kept as described in (38). A full description of the antimicrobial assay methods is in the supplementary information.

## Results and Discussion

### Hypothesis 1

#### Effect of environmental microbes and nutrients on oviposition

We tested whether *D. melanogaster* females plastically adjust egg number or clustering according to natural colonisation of the oviposition substrate by the microbes present in the environment. Within 48 hours of the oviposition assay, microbial growth was visible on 88% of substrates that lacked the antimicrobial preservatives propionic acid and Nipagin (Figure S2). In contrast, no microbial colonies were visible on substrates containing preservatives for the duration of the experiment. When preservatives were absent from the oviposition substrate, only 39.0% of groups laid any eggs, compared with 88.1% of groups laying when preservatives were present. Of the groups that did lay eggs, females laid significantly fewer when preservatives were absent (Figure 1A, Table S1-2, hurdle model with a negative binomial distribution, count part: *Z = 5.00*, *P* < 0.0001; binomial part: Z = 5.10, *P < 0.0001*), suggesting females are either sensitive to the absence of preservatives directly or to the increased presence of actively growing environmental microbes (the consequence of leaving out preservatives).

**Figure 1.**
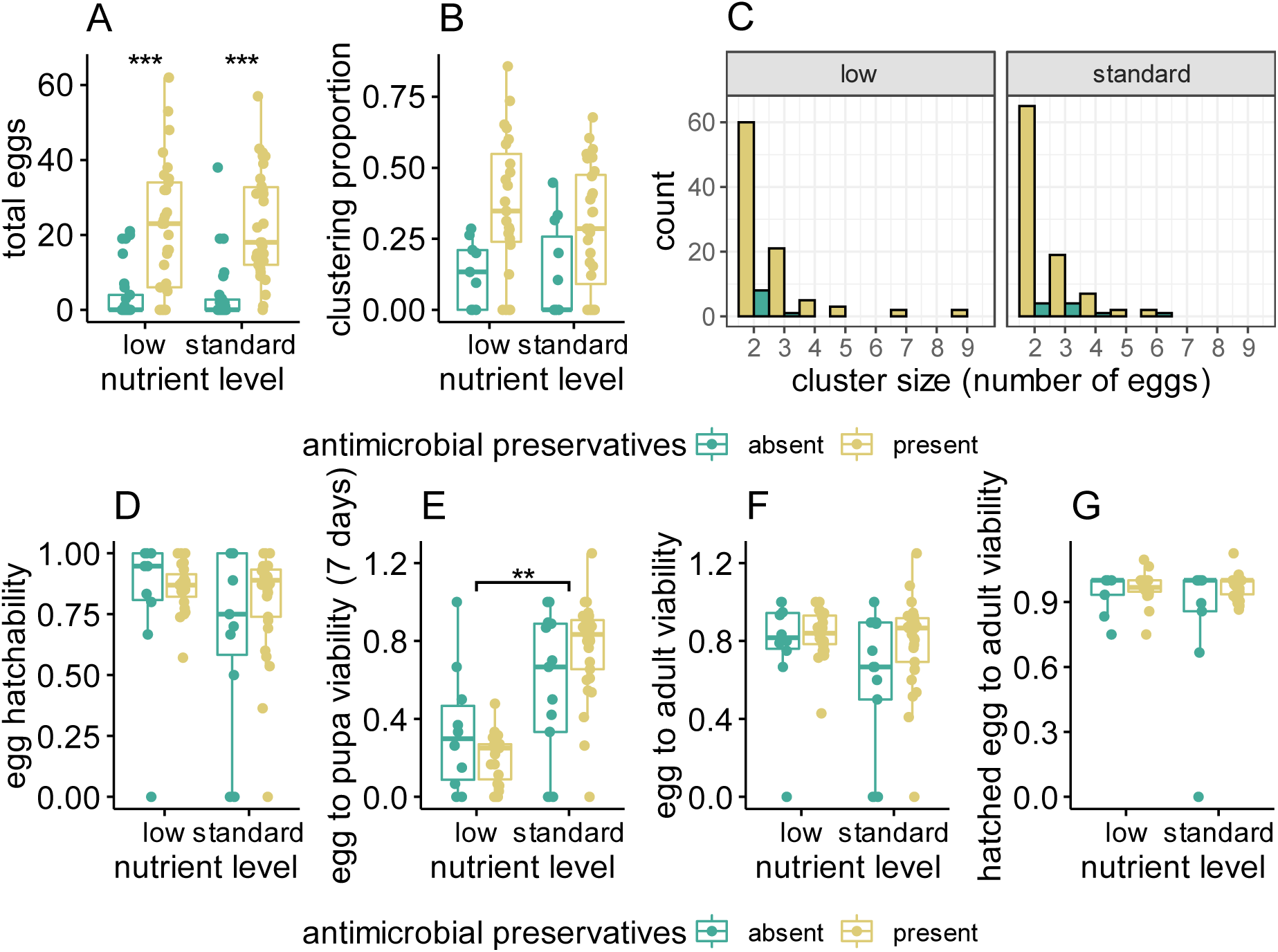
Females lay fewer eggs when antimicrobial preservatives are absent from the substrate. Oviposition substrates had low or standard levels of nutrients (yeast and sugar), and antimicrobial preservatives were either absent (green boxes and bars) or present (yellow boxes and bars). (A) total eggs laid by 4 females per oviposition vial. (B) the proportion of total eggs in each vial that were in any type of cluster (total clustered eggs / total eggs). (C) the number of egg clusters of different sizes, combined across all vials for each treatment. Bars are overlapping, not stacked. (D) the proportion of total eggs that had hatched after 48h (number of hatched eggs / total eggs) in each vial. (E) the proportion of total eggs which had developed into pupae after 7 days (number of pupae / total eggs) in each vial. (F) the proportion of total eggs that developed into adult offspring (total adult offspring / total eggs) in each vial. (G) the proportion of hatched eggs that developed into adult offspring (total adult offspring / number of hatched eggs) in each vial. Boxplots show the interquartile range (IQR) and median in the box, and whiskers represent the largest and smallest values within 1.5 times the IQR above and below the 75^th^ and 25^th^ percentiles, respectively. Raw data points are plotted with jitter. Statistically significant differences between treatments are indicated, using p values estimated from model testing (*** p < 0.0001, ** p <0.001).

The relationship between the microbial environment and *D. melanogaster* oviposition is likely to be complex since microbes can have beneficial, neutral and/or negative impacts on flies, depending on microbial species and their abundances. *D. melanogaster* oviposit into microbe-rich decomposing fruit, and indeed their larvae are dependent on beneficial yeasts for nutrition and normal development to adulthood. However, some bacteria and fungi (particularly moulds) are pathogenic when ingested by *D. melanogaster* larvae (25, 26). Although we did not characterise the species of microbes growing on the preservative-lacking substrates, we observed that most substrates harboured a mix of colony phenotypes, with several spore-bearing species characteristic of fungal moulds. Moulds such as *Penicillium* spp. are known to be detrimental to *D. melanogaster* development, likely because of the production of toxic secondary metabolites (39). A primary defence against being infected is for flies to avoid contact with harmful microorganisms, known as behavioural immunity (40). Consistent with this, detection of the microbial volatile geosmin leads to the suppression of feeding and egg-laying behaviours in *D. melanogaster* (39). It is possible that the flies in our experiments detected pathogenic microbes growing in the substrates lacking preservatives and either avoided contact with the substrate to protect themselves or retained their eggs to prevent infection of their offspring.

Although antimicrobial preservatives are added to artificial diets to control growth of mould and bacteria, they may simulate some of the microbial-derived metabolites that act as positive cues in natural oviposition sites. For example, yeast and bacteria produce short-chain fatty acids (SCFA) including propionic acid during fruit decomposition, and *Drosophila* possess neurons that are specifically activated by such acids (41). Indeed, female *D. melanogaster* adults exhibit attraction towards oviposition substrates containing SCFA (42). Similarly, ethanol (used to solubilise Nipagin) is one of the main metabolites of fermentation, and female *D. melanogaster* prefer to oviposit in ethanol-supplemented medium (43).

*Drosophila* oviposition preferences may also be affected by prior exposure to different diets. For example, prior exposure to normally repellent substances can reduce the strength of aversion through an apparent habituation effect (44). In the current study, females were housed on standard diet containing preservatives until the start of the experiment. Therefore, those females who were moved to the preservative-lacking substrates experienced a mismatch between the diets to which they had become habituated and the experimental substrate, the unfamiliarity of which may have contributed to females being less likely to lay eggs. However, nutrient level had no significant effect on the number of eggs laid (Figure 1A, Table S1-2), even though a similar mismatch in environment would apply to females moved from standard diet onto low nutrient substrates for the oviposition assay. We also conducted further experiments to test how oviposition is affected by the omission of different types of preservatives. These experiments, described below, revealed that specific preservatives are more likely than others to affect egg laying, weakening the idea that the novelty of an environment would be the sole cause of reduced fecundity.

Egg clustering proportions were calculated for all vials in which ≥2 eggs were laid (the minimum number of eggs required to form a cluster). The mean egg clustering proportion per vial was lower when preservatives were absent, but not significantly so (Figure 1B; Table S3). Clustering proportion increased significantly with the total number of eggs (F_(1,67)_ = 5.38, P = 0.02; Table S4, Figure S3), consistent with a pattern where females initially lay eggs singly and only later start to cluster eggs (10). This positive correlation between egg number and egg placement does not mean that females are clustering eggs by chance – indeed egg placement is predicted to be non-random, based on comparisons with null models simulating random placement (10), However, the relationship between egg number and egg clustering could explain why there was less clustering on substrates lacking preservatives, since fewer eggs were laid in the absence of preservatives. Egg cluster sizes ranged from 2-9, with the largest clusters found on low nutrient substrates (Figure 1C). Although the low nutrient substrates used in this experiment were not limiting for overall egg to adult viability, larvae took longer to develop into adults compared with flies reared on the standard nutrient diets (see below). Therefore, we might have expected females to reduce egg cluster sizes on low nutrient food to reduce competition between larvae. However, that larger clusters were laid on low nutrient substrates may indicate that larval cooperation could play a role when nutrients are scarce. *Drosophila* larvae are able to coordinate their feeding movements to feed more effectively (45) and larvae show greater aggregation on harder substrates, which are presumably more difficult to feed on (46). It remains to be investigated whether larvae emerging from clusters are better able to aggregate or coordinate feeding compared with larvae from eggs laid singly.

Eggs from the oviposition assay were scored for hatching, pupariation and eclosion. We excluded two vials from the hatching analysis because extensive microbial growth obscured hatching success. There were no significant effects of nutrient level, preservatives or clustering proportion on egg hatching success, although hatching was lowest on standard, preservative-free substrates (Figure 1D, Table S5-6). Standard nutrient substrates were more quickly covered in mould-type growth than low nutrient substrates (Figure S4), which could explain this lower (if not significant) hatching success. After 7 days, the proportion of laid eggs that had reached the pupal stage was significantly lower in the low nutrient treatments (F_(1,65)_ = 131.94, P < 0.0001), consistent with previous findings that lower nutrient levels increase development time (e.g. (47)). There was no significant effect of preservative presence on pupariation. However, there was a significant effect of the interaction between nutrient level and clustering proportion on pupariation (F_(1,65)_ = 5.07, P = 0.028) (Figure 1E, Table S9), likely driven by higher densities (and egg clustering) having a negative impact on development under low nutrient levels only. There were no significant effects of nutrient level, preservatives, total eggs or clustering proportion on egg-to-adult viability or on *hatched*-egg-to-adult viability (Figure 1F-G, Tables S10-S15). Moreover, hatched-egg-to-adult viability was generally high (mean 94.8%, Fig 1G), suggesting that egg hatching is the main hurdle for viability. It was surprising that egg-to-adult viability was unaffected by the extensive microbial growth in the absence of preservatives, given that in some cases the entire vial was swamped with mould which completely obscured the inside of the vial. It would be interesting to know whether there was a difference in the identity and pathogenicity of the microbial species growing in the vials in which females did lay eggs, compared with the vials in which they did not lay. Eggs that were laid mostly developed to adulthood successfully, suggesting that females made the correct decision to oviposit in those vials. However, we do not know whether the females who did not lay eggs also made the correct decision. A future experiment to test this could involve manually adding eggs to substrates that females have rejected as oviposition sites to test whether the eggs would have been viable had the females laid them there.

To test the extent to which females cluster their eggs with those of other females, and if such mixed maternity clustering is affected by substrate condition, we set up an experiment using low nutrient SYA ± preservatives in which we could distinguish the eggs and offspring of 1 focal female from those of 3 non-focals. There was significant agreement between the number of focal eggs we scored and the number of focal offspring that eclosed from each vial (ICC = 0.96, F_(59,60)_ = 55.1, p = 1.39e-36, Figure S5), meaning we were confident we could reliably distinguish the focal and non-focal eggs. Across all vials, there were 35 egg clusters containing ≥1 focal egg, and 25 of those clusters also contained ≥1 non-focal egg, meaning that focal eggs were part of a mixed maternity cluster in 71% of cases. Of the 35 clusters containing at least one focal egg, only 7 clusters were found in the no-preservative treatment, and, of these 7, only 2 were of mixed maternity (additional details in Figure S6). Given the low sample size in the no-preservative treatment, we lacked power to test the effect of substrate condition on the likelihood of mixed-maternity vs focal-only clustering. However, given mixed-maternity egg clustering was rarer in environments with higher microbial load, we found no support for our hypothesis that females would be more likely to lay eggs in clusters to benefit from ‘public good’ antimicrobial defences.

Overall, these experiments supported our hypothesis that females respond to differences in microbial environment by adjusting the number of eggs they lay. The following experiments were designed to separate out the effects on this phenomenon of the presence of actively growing microbes and the absence of preservatives, through alterations to the preservatives OR the microbial environment alone.

#### Effect of antimicrobial preservatives alone on oviposition

To uncouple the effects of microbial presence from the absence of the preservatives themselves, we compared oviposition on completely sterile substrates that only differed in the antimicrobial preservative added (propionic acid + Nipagin, propionic acid only, Nipagin (which is dissolved in ethanol), ethanol only, or no preservatives). In this experiment, all substrates were at standard nutrient levels since nutrient level had had no significant effect on fecundity in the earlier test (Figure 1). Overall, preservative treatment had a marginally significant effect on the number of eggs laid in each vial (Tables S16-17, Figure 2A). Compared with the standard treatment, which contained all preservatives, there were significantly fewer eggs laid when preservatives were completely absent, or when only Nipagin and/or ethanol were present (N + EtOH: Z = −2.53, P = 0.01; EtOH: Z = −2.01, P = 0.04; none: Z = - 2.61, P = 0.009). There were also fewer eggs laid on substates that contained propionic acid, but lacked Nipagin and ethanol, when compared to the standard treatment, but this difference was not statistically significant (Z = −1.41, P = 0.16). Combined, these results suggest that females are sensitive to antimicrobial preservatives when ovipositing, and it is the absence of propionic acid that had the largest effect on the number of eggs laid. Propionic acid is produced by bacteria during fermentation of rotting fruit – the natural site of *D. melanogaster* oviposition – and is detectable via specific olfactory receptors in the fly (41). Although adult flies have an aversion to propionic acid present at higher concentrations than used in the current study (2.5% vs 0.3% v/v) it is unknown whether lower concentrations would be as aversive, and propionic acid as an oviposition cue has not been investigated (41). In contrast to adults, *D. melanogaster* larvae are attracted to propionic acid, and supplementation of nutrient-poor media with 1% propionic acid can improve larval survival (48). It is therefore possible that females increase egg laying at certain concentrations of propionic acid since it represents a good developmental environment for their offspring. This may partly explain the reduced egg laying seen on substrates lacking preservatives in the first experiment, but absence of propionic acid alone is unlikely to explain the high number of vials with zero eggs seen in the first, but not the second, experiment. The key difference between the two experiments was the sterility of oviposition substrates upon exposure to a female. This suggests that some females refrained from laying in the first experiment due to the microbial environment, rather than as a response to the absence of preservatives *per se*.

**Figure 2.**
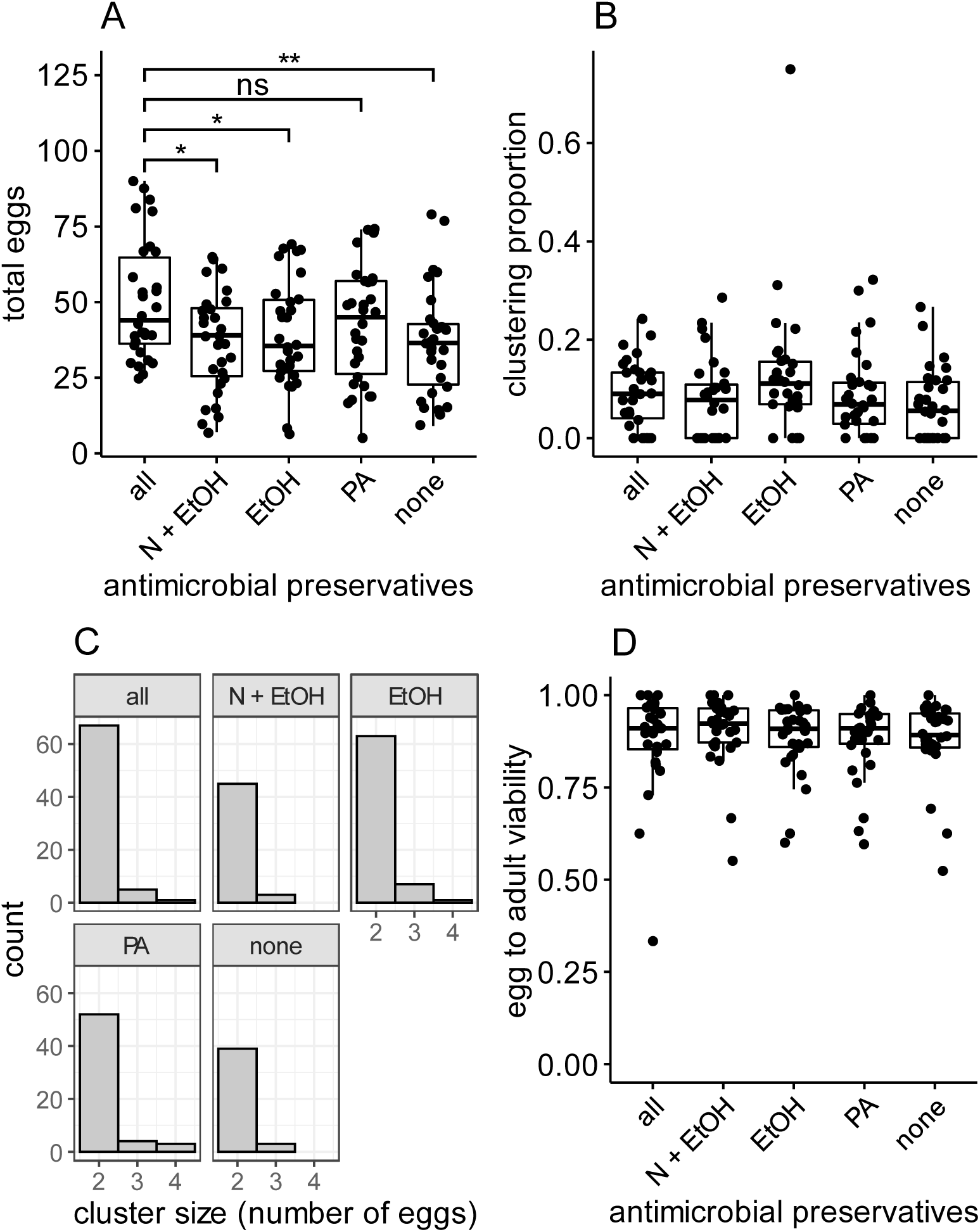
Antimicrobial preservatives have a marginally significant effect on fecundity. Oviposition substrates contained Nipagin and propionic acid (“all”), Nipagin (“N + EtOH”), Ethanol only (“EtOH”), propionic acid only (“PA”) or no preservatives (“none”). (A) total eggs laid by 4 females per oviposition vial. (B) the proportion of total eggs in each vial that were in any type of cluster (total clustered eggs / total eggs). (C) the number of egg clusters of different sizes, combined across all vials for each treatment. (D) the proportion of total eggs that developed into adult offspring (total adult offspring / total eggs) in each vial. Boxplots are as decribed for Figure 1. Statistical significance indicated in (A) (* p < 0.01; ** p < 0.001; ns: p > 0.05) with p values derived from model summary.

Overall, the egg clustering proportion was lower than in the first experiment (Figure 2B, Table S18), despite the total number of eggs being higher. There was no significant effect of treatment on clustering proportion (F_(4, 144)_ = 1.57, P = 0.19), but clustering proportion significantly increased with the number of eggs laid (F_(1,144)_, P < 0.0001, Table S19, Figure S7). Therefore, although females adjusted the number of eggs they laid when different preservatives were present, they did not adjust how they placed those eggs. Egg to adult viability was not significantly affected by treatment, total eggs or clustering proportion (Figure 2D, Tables S20-22).

#### Effect of commensal and pathogenic microbes on oviposition

The first experiment utilised environmental microbes that naturally colonised the non-sterile substrates which lacked preservatives. The visible microbial growth on these substrates appeared to mostly be fungal moulds. To test instead for the effects of fly-associated microbial communities and known pathogens on oviposition, we inoculated the surface of sterile oviposition substrates (which all lacked preservatives) with either commensal microbes, the reportedly entomopathogenic bacterium *Alcaligenes faecalis* (25), or sterile control washes. We predicted that females would cluster their eggs more when pathogenic microbes were present when compared with commensal microbes if egg clustering provides benefits from public antimicrobial defences. To verify that the washes lead to differences in microbial environment, we checked the substrates 48 h following oviposition for visible microbial colonies. There were visible colonies in 90% of the commensal microbe substrates, compared with 27% of negative control substrates, 20% of fly background control substrates and 23% of *A. faecalis* substrates. We did not investigate the species identity of any microbial colonies, but the colonies visible in the *A. faecalis* treatment did not have the morphology of *A. faecalis,* suggesting this bacterium does not grow as quickly as other species, or at all, on SYA media. In a subset of substrates that remained unexposed to live flies, there was visible microbial growth on 5 out of 6 commensal wash substrates, and 1 out of 5 negative control substrates, but no colonies were visible on any of the fly background control or *A. faecalis* substrates. Combined, these observations showed that, as intended, microbes were successfully transferred to the oviposition substrates in the commensal microbe wash, but not the negative or background controls.

Despite the established differences in microbial environment across vials, there was no significant effect of substrate treatment on the number of eggs laid, or the egg clustering proportion (Table S23-26, Figure 3A-B) although clustering proportion was again significantly affected by the total number of eggs (F_(1,110)_ = 7.9, P = 0.006, Table S26, Figure S8). Therefore, we found no support for the hypothesis that flies increase egg clustering in response to pathogens. However, it could be that different strains of *A. faecalis* differ in their entomopathogenicity. A previous study of *A. faecalis* pathogenicity reported a 25% mortality rate upon larval ingestion (25), which was not seen in our experiment - egg to adult viability was unaffected by substrate treatment and remained high at 88% despite extensive microbial growth in many vials (Tables S27-29, Figure 3D). It is therefore possible that females did not alter their oviposition in response to the presence of this strain of *A. faecalis* due to insufficient cues of a pathogenic environment. It is also unknown whether *A. faecalis* produces volatiles that could be detected and cause aversion in fruit flies, akin to geosmin produced by some microbes. Additionally, since no *A. faecalis* colonies were visible on the substrate surface, it is possible the culture was not actively growing, or growing very slowly, which could reduce the probability of its detection by females. Experiments using multiple verified entomopathogenic species would be necessary to further investigate whether such microbes can affect oviposition decisions. Interestingly, oviposition was also unaffected by the commensal microbe treatment. The diversity of commensal microbes (which should contain the transient gut microbiota of flies) is likely to be distinct from the environmental microbes that would have colonised the substrates in the absence of preservatives in the first experiment. There is evidence that *Drosophila* can distinguish between commensal and pathogenic microbes and select commensal-rich sites for egg-laying (23). The commensal microbial community can produce anti-fungal metabolites as well as provide access to nutrients which supports larval development (49, 50). These beneficial properties of a commensal microbial community may explain why females were not averse to laying eggs in this assay.

**Figure 3.**
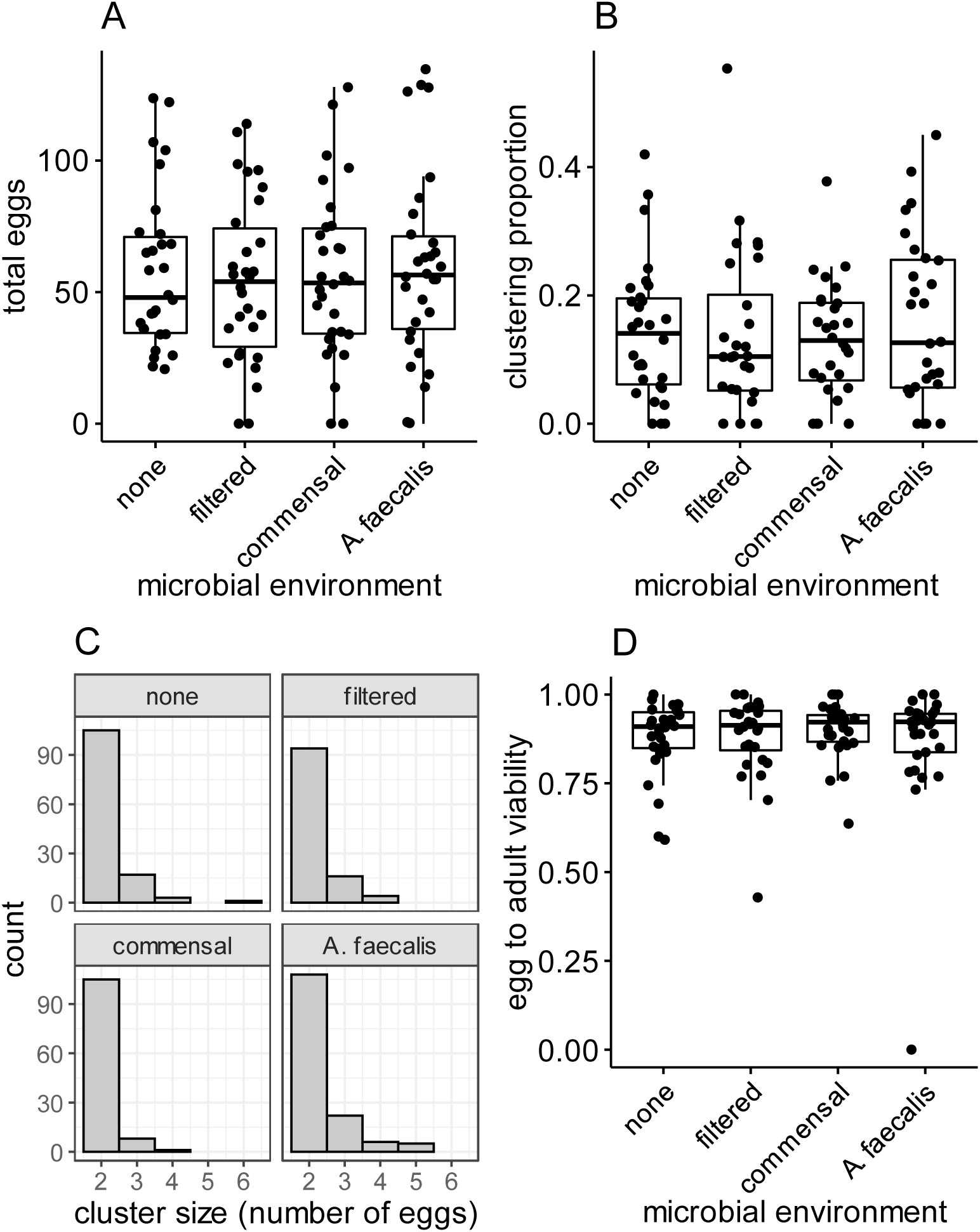
Neither the presence of commensal microbes or the bacterium *Alcaligenes faecalis* affects oviposition or offspring viability. Oviposition substrates lacking preservatives were seeded with one of four different washes: negative control (“none”), fly background control (“filtered”), fly commensal microbes (“commensal”), and a culture of the bacterium *Alcaligenes faecalis* M3A (“A. faecalis”). (A) total eggs laid by 4 females per oviposition vial. (B) the proportion of total eggs in each vial that were in any type of cluster (total clustered eggs / total eggs). (C) the number of egg clusters of different sizes, combined across all vials for each treatment. (D) the proportion of total eggs that developed into adult offspring (total adult offspring / total eggs) in each vial. Boxplots are as decribed for Figure 1.

### Hypothesis 2

#### Antimicrobial activity of egg surface molecules

In the final set of experiments, we tested if *Drosophila melanogaster* eggs or laid egg soluble material (LESM) exhibit antimicrobial activity, by conducting antimicrobial peptide diffusion assays against four species of microbes – the gram-negative strains *Escherichia coli* DH5α and *Alcaligenes faecalis* M3A, the yeast *Saccharomyces cerevisiae*, and the gram-positive bacteria *Micrococcus luteus*. We also included eggs and egg wash from the Medfly *C. capitata* as a positive control (15). There were clear zones of growth inhibition around wells containing Medfly LESM for all four species of microbes, but *D. melanogaster* LESM exhibited no antimicrobial activity (Figure 4). Additionally, we quantified the total protein amount in each egg wash using a Qubit protein assay. For the Medfly sample, the protein concentration was 834 µg/ml. We therefore calculated the protein amount per egg to be 83 ng. For *D. melanogaster*, the protein concentration was below the limit of detection for the Qubit. Therefore, counter to our hypothesis, we found no evidence that *D. melanogaster* females provisioned their eggs with soluble peptides, and found no evidence for any broad range antimicrobial activity on the egg surface as occurs in Medfly (15). Whole eggs from *D. melanogaster* also did not exhibit any antimicrobial activity when tested against *E. coli* (Figure S9). Overall, we found no evidence to support the hypothesis that *D. melanogaster* females provision their eggs with antimicrobials. Despite the evidence that AMP genes are expressed and enriched for expression in the female reproductive tract, none of the 21 known AMPs, or the 12 Bomanin peptides (51) were found among the 1840 proteins identified in a recent proteomic study of the female reproductive tissue and luminal fluid (52). It is possible that AMP genes are not translated at high levels in the female reproductive tract, or that AMPs are produced under specific conditions that were not used in the proteomics study. Regardless, the absence of antimicrobial activity in our diffusion assays suggests *D. melanogaster* do not provision their eggs with antimicrobial defences that could be exploited as public goods. Since Drosophila lay into microbially-rich environments and are dependent on microbial phytophagous activity to break down fruits and provide nutrients to the flies, it is possible that deploying broad-acting antimicrobials on egg surfaces is detrimental if doing so depletes some of the beneficial microbial species. Instead, it could be that *D. melanogaster* protect their offspring from infection by avoiding ovipositing into sites containing pathogens (behavioural immunity) or choosing sites where the microbial community itself is producing antimicrobials against entomopathogenic species (49).

**Figure 4.**
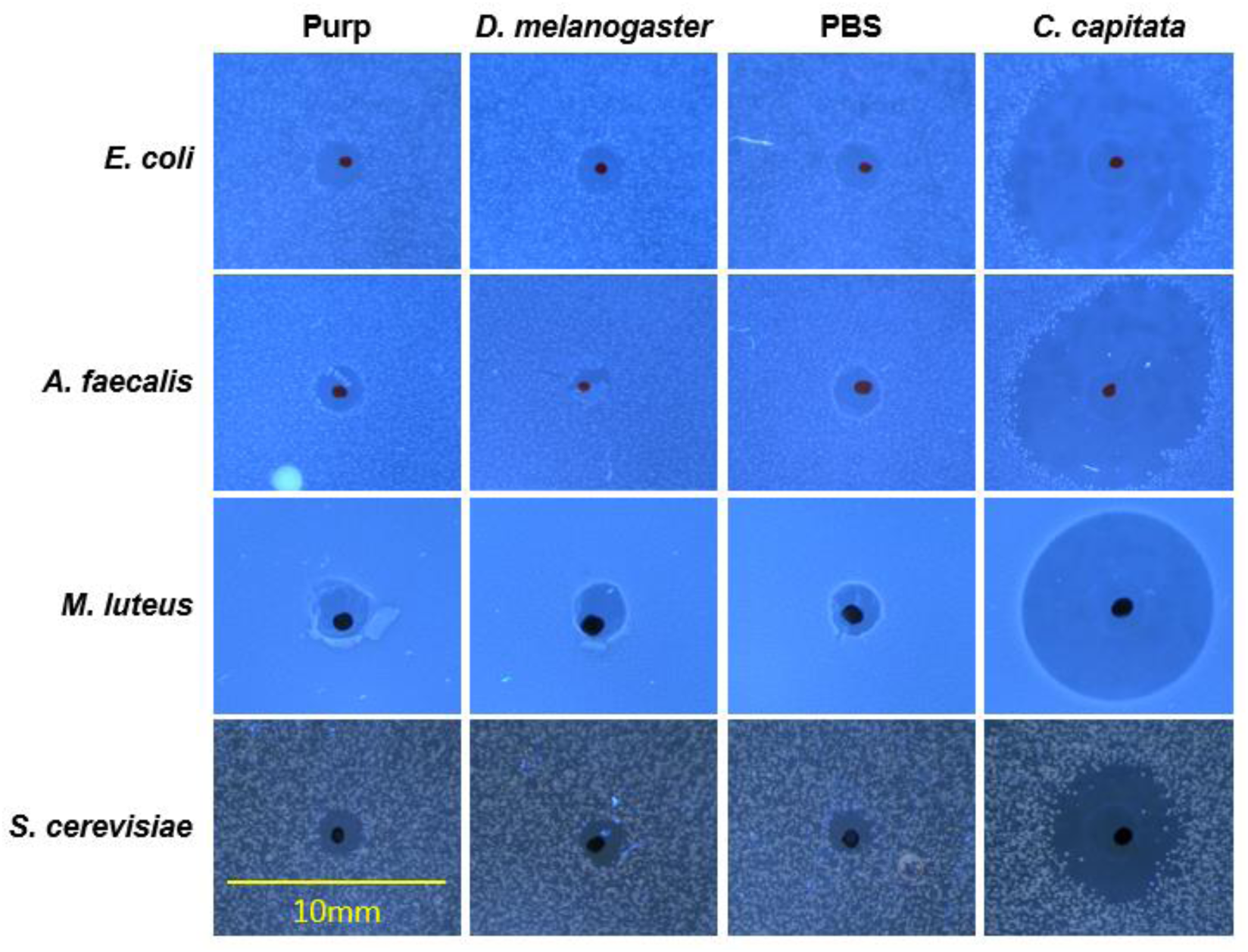
Laid egg soluble material of *Drosophila melanogaster* does not exhibit broad-acting antimicrobial activity. Soluble material from washing freshly laid *D. melanogaster* or *C. capitata* eggs was pipetted directly into wells in the assay plates. Negative controls were PBS only for *C. capitata* eggs (column 3) and PBS that had been in contact with to a fragment of purple grape juice agar for *D. melanogaster* eggs (column 1, “purp”). Each plate contained a live culture of either *Escherichia coli* dh5α, *Alcaligenes faecalis* M3A, *Micrococcus luteus* or *Saccharomyces cerevisiae* NYCC 505. Individual wells were photographed under a microscope to show any zones of growth inhibition surrounding the well, and these images were arranged in the above figure. The centre of each well was marked with a black dot on the petri dish for ease of identification. A 10 mm scale bar is shown at the bottom left.

### Conclusions

Our first hypothesis (*D. melanogaster* plastically adjust egg placement decisions based on the microbial environment) received partial support. We found that females reduced egg laying on substrates lacking preservatives with environmentally-derived microbes, but they did not significantly change the extent to which eggs were clustered. Females did not adjust their egg laying in the presence of commensal microbes, or the gram negative species *A. faecalis,* implying that different microbial environments elicit different oviposition responses in female fruit flies. It is possible that the environmentally derived (i.e. non-commensal) microbial community contained pathogenic species, to which the females responded by reducing or abstaining from egg laying. Further work is required to better characterise the microbial environments and how this relates to the oviposition decisions of females. Related to our first hypothesis, we also found no evidence of an increase in mixed maternity egg clustering in the presence of environmentally-derived microbes.

Our second hypothesis (*D. melanogaster* eggs exhibit broad spectrum antimicrobial activity) was not supported. We found no antimicrobial effects of *D. melanogaster* of the soluble material from the surface of eggs, or of the whole eggs themselves. We cannot rule out the possibility of species-specific antimicrobial compounds being present as only 4 microbial species were tested, but the *D. melanogaster* eggs certainly did not exhibit the same type of broad-spectrum antimicrobial activity as Medfly eggs. This finding, combined with the fact females do not adjust egg clustering, or increase mixed maternity clustering, in the presence of microbes suggests *D. melanogaster* females do not cluster their eggs to gain public goods benefits from the communal production of antimicrobial compounds.

## Acknowledgements

We would like to thank Paul Candon and Kerri Armstrong for technical assistance.

