## Supplementary information for "Female oviposition decisions are influenced by the microbial environment"

Fowler, EK., Friend, LA., Churchill, ER., Yu, DW., Archetti, M., Bourke, AFG., Bretman, A., and Chapman, T.

### Supplementary Information

#### Supplementary Methods

*Antimicrobial activity of egg surface molecules*

To test if *D. melanogaster* eggs exhibit antimicrobial activity, we conducted radial diffusion assays according to (25). The bacteria *Escherichia coli* dh5α, *Alcaligenes faecalis* M3A and *Micrococcus luteus* were grown overnight in Lysogeny Broth (LB: 5g NaCl, 10 g tryptone, 5 g yeast extract and 1.5 g glucose per litre $H_2O$) and the yeast *Saccharomyces cerevisiae* NYCC 505 was grown overnight in YPD medium (10 g yeast extract, 20 g peptone and 20 g glucose per litre $H_2O$). Overnight cultures were inoculated 1:100 into fresh broth and grown for 4 hours at 180 RPM and 30°C. 50 ml of each culture was centrifuged at 4°C for 10 mins, washed in 10 ml ice cold 10 mM sodium phosphate buffer, centrifuged once more and finally resuspended in 5 ml ice cold sodium phosphate buffer.

The CFU/ml of the resuspended microbial cultures was estimated from OD600 measurements and the volume equivalent to $2.54 \times 10^6$ CFU was used to inoculate the underlay agarose medium in the diffusion assay. The underlay medium for *E. coli*, *M. luteus* and *A. faecalis* consisted of 50 ml 100 mM sodium phosphate, 5 ml standard LB (recipe above), 5 g agarose and 445 ml H2O. The underlay medium for *S. cerevisiae* was identical, but with 5 ml standard YPD broth in place of LB. Once autoclaved and cooled to 42°C, 7 ml of the underlay medium was mixed with ~$2.54 \times 10^6$ CFU of the focal microbial species and poured into 90 mm petri dishes (*Fisher Scientific* #12654785) to make a very thin layer. Once set, a modified sterile p1000 Gilson pipette tip was used to punch four holes of ~2 mm in the underlay. One hole was punched for each of the samples being tested (two negative controls, *Drosophila* egg wash, and Medfly egg wash).

To generate the laid egg soluble material (LESM), we allowed *D. melanogaster* females to lay onto purple grape juice agar plates before picking 1000 eggs to 50 µl of PBS. For Medfly, eggs were collected dry as females pushed them through a mesh onto a piece of foil. 500 Medfly eggs were transferred to 50 µl of PBS. Since purple grape juice agar plates contain Nipagen, a negative control for the *D. melanogaster* eggs was made by transferring a small piece of grape juice agar to 50 µl PBS, to control for any small amount of grape juice agar that may have been transferred with the eggs. After incubating the eggs or grape juice agar in PBS for 5 minutes, we centrifuged the samples to pellet the insoluble material and collected the supernatant. The supernatant (LESM) was passed through a spin filter before use in the assays (*Corning Costar* #8163).

We quantified the amount of protein in each LESM wash using a Qubit assay according to kit protocol, and then 2 µl of the LESM samples, or negative controls (PBS for Medfly eggs, PBS exposed to purple grape juice agar for *D. melanogaster* eggs) were applied to each of the wells of the underlay medium. The underlay was then incubated at 30°C for 3 hours. The overlay gel consisted of an enriched nutrient agarose. For *E. coli*, *M. luteus* and *A. faecalis* the overlay medium was made to the following recipe: 5 g NaCl, 10 g tryptone, 5 g yeast extract, 1.5 g glucose, 5 g agarose and 500 ml RO $H_2O$, and for *S. cerevisiae* the overlay medium was 10 g yeast extract, 20 g peptone, 20 g glucose, 5 g agarose and 500 ml $H_2O$. Once autoclaved and cooled to 42°C, 8 ml of overlay agarose was poured over the underlay. Plates were then incubated at 30°C overnight. The following day, plates were photographed using a GXCAM HiChrome-S camera (*GT Vision Ltd.*) mounted on a Leica MZ75 dissecting microscope.

### Supplementary Tables

**Table S1.** Summary statistics for total eggs laid on substrates of different nutrient level and preservative presence.

| nutrient | preservatives | N | mean total eggs | sd | se | ci |
|---|---|---|---|---|---|---|
| low | absent | 29 | 3.97 | 7.18 | 1.33 | 2.73 |
| low | present | 29 | 21.72 | 17.45 | 3.24 | 6.64 |
| standard | absent | 30 | 3.77 | 8.24 | 1.50 | 3.08 |
| standard | present | 30 | 22.53 | 14.44 | 2.64 | 5.39 |

**Table S2.** Effect of nutrient level and preservative presence on number of eggs laid. Model coefficients of a two part hurdle model.

| **Model formula** TOTAL.EGGS~NUTRIENT + PRESERVATIVES, data = EXP1A.DATA, dist = “negbin” | | | | |
|---|---|---|---|---|
| **Count model coefficients (truncated negbin with log link)** | | | | |
| **Variable** | **Estimate** | **Std. Error** | **z value** | **Pr(>\|z\|)** |
| (Intercept) | 2.3689 | 0.1874 | 12.643 | < 2e-16 *** |
| NUTRIENTstandard | -0.1997 | 0.1729 | -1.155 | 0.247935 |
| PRESERVATIVESpresent | 0.9575 | 0.1917 | 4.995 | 5.87e-07 *** |
| Log(theta) | 0.7144 | 0.2050 | 3.485 | 0.000491 *** |
| **Zero hurdle model coefficients (binomial with logit link)** | | | | |
| **Variable** | **Estimate** | **Std. Error** | **z value** | **Pr(>\|z\|)** |
| (Intercept) | -0.8622 | 0.3718 | -2.319 | 0.0204 * |
| NUTRIENTstandard | 0.7808 | 0.4588 | 1.702 | 0.0888 . |
| PRESERVATIVESpresent | 2.5281 | 0.4961 | 5.096 | 3.46e-07 *** |
| Log-likelihood: -347.3 on 7 Df | | | | |

**Table S3.** Summary statistics for clustering proportion on substrates of different nutrient level and preservative presence.

| nutrient | preservatives | N | mean clustering proportion | sd | se | ci |
|---|---|---|---|---|---|---|
| low | absent | 9 | 0.13 | 0.11 | 0.04 | 0.09 |
| low | present | 23 | 0.37 | 0.27 | 0.05 | 0.11 |
| standard | absent | 11 | 0.18 | 0.17 | 0.05 | 0.11 |
| standard | present | 28 | 0.29 | 0.22 | 0.04 | 0.08 |

**Table S4.** Effect of nutrient level, preservative presence and total eggs on the proportion of eggs in clusters. Model coefficients of a quasibinomial GLM.

<table>
<tr><td colspan="5">Model formula<br>y1 ~ NUTRIENT + PRESERVATIVES + TOTAL.EGGS, family = quasibinomial, data = EXP1A.DATA.M1B</td></tr>
<tr><td colspan="5">Model coefficients (quasibinomial GLM)</td></tr>
<tr><td>Variable</td><td>Estimate</td><td>Std. Error</td><td>t value</td><td>Pr(>|t|)</td></tr>
<tr><td>(Intercept)</td><td>-1.583427</td><td>0.351746</td><td>-4.502</td><td>2.76e-05 ***</td></tr>
<tr><td>NUTRIENTstandard</td><td>-0.060146</td><td>0.200869</td><td>-0.299</td><td>0.7655</td></tr>
<tr><td>PRESERVATIVESpresent</td><td>0.518383</td><td>0.333448</td><td>1.555</td><td>0.1247</td></tr>
<tr><td>TOTAL.EGGS</td><td>0.017107</td><td>0.007422</td><td>2.305</td><td>0.0243 *</td></tr>
<tr><td colspan="5">(Dispersion parameter for quasibinomial family taken to be 3.390113)<br>Null deviance: 306.14 on 70 degrees of freedom<br>Residual deviance: 265.22 on 67 degrees of freedom</td></tr>
<tr><td colspan="5">Analysis of Deviance Table (Type II tests)</td></tr>
<tr><td>Variable</td><td>Sum Sq</td><td>Df</td><td>F value</td><td>P value</td></tr>
<tr><td>NUTRIENT</td><td>0.304</td><td>1</td><td>0.0896</td><td>0.76556</td></tr>
<tr><td>PRESERVATIVES</td><td>8.644</td><td>1</td><td>2.5498</td><td>0.11501</td></tr>
<tr><td>TOTAL.EGGS</td><td>18.229</td><td>1</td><td>5.3770</td><td>0.02346 *</td></tr>
<tr><td>residuals</td><td>227.137</td><td>67</td><td></td><td></td></tr>
</table>

**Table S5.** Summary statistics for hatching success of eggs laid onto substrates of different nutrient level and preservative presence.

| **nutrient** | **preservatives** | **N** | **mean egg hatchability** | **sd** | **se** | **ci** |
|---|---|---|---|---|---|---|
| low | absent | 10 | 0.82 | 0.31 | 0.10 | 0.22 |
| low | present | 23 | 0.86 | 0.10 | 0.02 | 0.04 |
| standard | absent | 11 | 0.68 | 0.38 | 0.11 | 0.25 |
| standard | present | 29 | 0.81 | 0.22 | 0.04 | 0.08 |

**Table S6.** Effect of nutrient level, preservative presence and clustering proportion on egg hatchability. Model coefficients of a quasibinomial GLM.

| **Model formula**<br>Y2 ~ NUTRIENT + PRESERVATIVES + PROPORTION.CLUSTERED, family = quasibinomial, data = EXP1A.DATA.M1C | | | | |
|---|---|---|---|---|
| **Model coefficients (quasibinomial GLM)** | | | | |
| **Variable** | **Estimate** | **Std. Error** | **t value** | **Pr(>\|t\|)** |
| (Intercept) | 1.2110 | 0.3887 | 3.116 | 0.00273 ** |
| NUTRIENTstandard | -0.1189 | 0.2718 | -0.437 | 0.66323 |
| PRESERVATIVESpresent | 0.3193 | 0.4210 | 0.758 | 0.45093 |
| PROPORTION.CLUSTERED | 0.3783 | 0.7291 | 0.519 | 0.60561 |
| (Dispersion parameter for quasibinomial family taken to be 3.756746)<br>Null deviance: 252.30 on 68 degrees of freedom<br>Residual deviance: 246.93 on 65 degrees of freedom | | | | |
| **Analysis of Deviance Table (Type II tests)** | | | | |
| **Variable** | **Sum Sq** | **Df** | **F value** | **P value** |
| NUTRIENT | 0.72 | 1 | 0.1915 | 0.6631 |
| PRESERVATIVES | 2.10 | 1 | 0.5591 | 0.4573 |
| PROPORTION.CLUSTERED | 1.01 | 1 | 0.2687 | 0.6059 |
| residuals | 244.19 | 65 | | |

**Table S7.** Summary statistics for egg to pupa viability, 7 days post-oviposition, for eggs laid on substrates of different nutrient level and preservative presence.

| **nutrient** | **preservatives** | **N** | **mean egg to pupa viability** | **sd** | **se** | **ci** |
|---|---|---|---|---|---|---|
| low | absent | 10 | 0.33 | 0.32 | 0.10 | 0.23 |
| low | present | 23 | 0.20 | 0.13 | 0.03 | 0.05 |
| standard | absent | 13 | 0.56 | 0.38 | 0.11 | 0.23 |
| standard | present | 28 | 0.75 | 0.23 | 0.04 | 0.09 |

**Table S8.** Effect of nutrient level, preservative presence and total eggs on egg to pupa viability. Model coefficients of a quasibinomial GLM.

| **Model formula** | | | | |
|---|---|---|---|---|
| Y3 ~ NUTRIENT + PRESERVATIVES + TOTAL.EGGS, family = quasibinomial, data = EXP1.DATA.M1D) | | | | |
| **Model coefficients (quasibinomial GLM)** | | | | |
| **Variable** | **Estimate** | **Std. Error** | **t value** | **Pr(>\|t\|)** |
| (Intercept) | -1.038283 | 0.350161 | -2.965 | 0.00413 ** |
| NUTRIENTstandard | 2.311236 | 0.224338 | 10.302 | 1.14e-15 *** |
| PRESERVATIVESpresent | 0.206939 | 0.331148 | 0.625 | 0.53406 |
| TOTAL.EGGS | -0.014202 | 0.008449 | -1.681 | 0.09724 . |
| (Dispersion parameter for quasibinomial family taken to be 3.434935)<br>Null deviance: 710.45 on 73 degrees of freedom<br>Residual deviance: 261.75 on 70 degrees of freedom | | | | |
| **Analysis of Deviance Table (Type II tests)** | | | | |
| **Variable** | **Sum Sq** | **Df** | **F value** | **P value** |
| NUTRIENT | 428.56 | 1 | 124.7667 | < 2e-16 *** |
| PRESERVATIVES | 1.34 | 1 | 0.3900 | 0.5343 |
| TOTAL.EGGS | 9.85 | 1 | 2.8662 | 0.0949 . |
| residuals | 240.44 | 70 | | |

**Table S9.** Effect of nutrient level, preservative presence and clustering proportion on egg to pupa viability. Model coefficients of a quasibinomial GLM.

| **Model formula**<br>Y3 ~ NUTRIENT + PRESERVATIVES + PROPORTION.CLUSTERED + NUTRIENT:PROPORTION.CLUSTERED, family = quasibinomial, data = EXP1A.DATA.M1D | | | | |
|---|---|---|---|---|
| **Model coefficients (quasibinomial GLM)** | | | | |
| **Variable** | **Estimate** | **Std. Error** | **t value** | **Pr(>\|t\|)** |
| (Intercept) | -0.97408 | 0.38121 | -2.555 | 0.01296 * |
| NUTRIENTstandard | 1.46123 | 0.44757 | 3.265 | 0.00175 ** |
| PRESERVATIVESpresent | 0.04433 | 0.31950 | 0.139 | 0.89007 |
| PROPORTION.CLUSTERED | -1.09096 | 0.90528 | -1.205 | 0.23253 |
| NUTRIENTstandard: PROPORTION.CLUSTERED | 2.62547 | 1.17547 | 2.234 | 0.02896 * |
| (Dispersion parameter for quasibinomial family taken to be 3.380971)<br>Null deviance: 705.05 on 69 degrees of freedom<br>Residual deviance: 241.70 on 65 degrees of freedom | | | | |
| **Analysis of Deviance Table (Type II tests)** | | | | |
| **Variable** | **Sum Sq** | **Df** | **F value** | **P value** |
| NUTRIENT | 446.10 | 1 | 131.9449 | < 2e-16 *** |
| PRESERVATIVES | 0.07 | 1 | 0.0192 | 0.89013 |
| PROPORTION.CLUSTERED | 1.67 | 1 | 0.4940 | 0.48467 |
| NUTRIENT: PROPORTION.CLUSTERED | 17.15 | 1 | 5.0737 | 0.02767 * |
| residuals | 219.76 | 65 | | |

**Table S10.** Summary statistics for egg to adult viability, for eggs laid on substrates of different nutrient level and preservative presence.

| **nutrient** | **preservatives** | **N** | **mean egg to adult viability** | **sd** | **se** | **ci** |
|---|---|---|---|---|---|---|
| low | absent | 10 | 0.77 | 0.29 | 0.09 | 0.21 |
| low | present | 23 | 0.84 | 0.12 | 0.03 | 0.05 |
| standard | absent | 13 | 0.60 | 0.37 | 0.10 | 0.22 |
| standard | present | 27 | 0.77 | 0.21 | 0.04 | 0.08 |

**Table S11.** Effect of nutrient level, preservative presence and total eggs on egg to adult viability. Model coefficients of a quasibinomial GLM.

| **Model formula** | | | | |
|---|---|---|---|---|
| Y4 ~ NUTRIENT + PRESERVATIVES + TOTAL.EGGS, family = quasibinomial, data = EXP1A.DATA.M1E | | | | |
| **Model coefficients (quasibinomial GLM)** | | | | |
| **Variable** | **Estimate** | **Std. Error** | **t value** | **Pr(>\|t\|)** |
| (Intercept) | 1.282732 | 0.361394 | 3.549 | 0.0007 *** |
| NUTRIENTstandard | -0.126730 | 0.238887 | -0.531 | 0.5975 |
| PRESERVATIVESpresent | 0.323472 | 0.343421 | 0.942 | 0.3495 |
| TOTAL.EGGS | -0.003835 | 0.008953 | -0.428 | 0.6697 |
| (Dispersion parameter for quasibinomial family taken to be 3.464615)<br>Null deviance: 237.54 on 72 degrees of freedom<br>Residual deviance: 233.68 on 69 degrees of freedom | | | | |
| **Analysis of Deviance Table (Type II tests)** | | | | |
| **Variable** | **Sum Sq** | **Df** | **F value** | **P value** |
| NUTRIENT | 0.976 | 1 | 0.2819 | 0.5972 |
| PRESERVATIVES | 2.995 | 1 | 0.8645 | 0.3557 |
| TOTAL.EGGS | 0.635 | 1 | 0.1833 | 0.6699 |
| residuals | 239.046 | 69 | | |

**Table S12.** Effect of nutrient level, preservative presence and clustering proportion on egg to adult viability. Model coefficients of a quasibinomial GLM.

| **Model formula** | | | | |
|---|---|---|---|---|
| Y4 ~ NUTRIENT + PRESERVATIVES + PROPORTION.CLUSTERED, family = quasibinomial, data = EXP1A.DATA.M1E | | | | |
| **Model coefficients (quasibinomial GLM)** | | | | |
| **Variable** | **Estimate** | **Std. Error** | **t value** | **Pr(>\|t\|)** |
| (Intercept) | 1.10937 | 0.34498 | 3.216 | 0.00203 ** |
| NUTRIENTstandard | -0.09174 | 0.23806 | -0.385 | 0.70124 |
| PRESERVATIVESpresent | 0.15766 | 0.33366 | 0.473 | 0.63814 |
| PROPORTION.CLUSTERED | 0.52538 | 0.64598 | 0.813 | 0.41901 |
| (Dispersion parameter for quasibinomial family taken to be 3.425762)<br>Null deviance: 227.50 on 68 degrees of freedom<br>Residual deviance: 222.79 on 65 degrees of freedom | | | | |
| **Analysis of Deviance Table (Type II tests)** | | | | |
| **Variable** | **Sum Sq** | **Df** | **F value** | **P value** |
| NUTRIENT | 0.509 | 1 | 0.1486 | 0.7011 |
| PRESERVATIVES | 0.753 | 1 | 0.2198 | 0.6407 |
| PROPORTION.CLUSTERED | 2.264 | 1 | 0.6609 | 0.4192 |
| residuals | 222.675 | 65 | | |

**Table S13.** Summary statistics for hatched egg to adult viability for eggs laid on substrates of different nutrient level and preservative presence.

| nutrient | preservatives | N | mean hatched egg to adult viability | sd | se | ci |
|---|---|---|---|---|---|---|
| low | absent | 9 | 0.95 | 0.09 | 0.03 | 0.07 |
| low | present | 21 | 0.96 | 0.06 | 0.01 | 0.03 |
| standard | absent | 9 | 0.82 | 0.33 | 0.11 | 0.25 |
| standard | present | 24 | 0.96 | 0.04 | 0.01 | 0.02 |

**Table S14.** Effect of nutrient level, preservative presence and total eggs on hatched egg to adult viability. Model coefficients of a quasibinomial GLM.

<table>
<tr><td colspan="5">Model formula<br>Y5 ~ NUTRIENT + PRESERVATIVES + TOTAL.EGGS, family = quasibinomial, data = EXP1A.DATA.M1F</td></tr>
<tr><td colspan="5">Model coefficients (quasibinomial GLM)</td></tr>
<tr><td>Variable</td><td>Estimate</td><td>Std. Error</td><td>t value</td><td>Pr(>|t|)</td></tr>
<tr><td>(Intercept)</td><td>2.649327</td><td>0.430491</td><td>6.154</td><td>7.16e-08 ***</td></tr>
<tr><td>NUTRIENT standard</td><td>-0.219312</td><td>0.327478</td><td>-0.670</td><td>0.506</td></tr>
<tr><td>PRESERVATIVES present</td><td>0.775629</td><td>0.480758</td><td>1.613</td><td>0.112</td></tr>
<tr><td>TOTAL.EGGS</td><td>-0.004746</td><td>0.012258</td><td>-0.387</td><td>0.700</td></tr>
<tr><td colspan="5">(Dispersion parameter for quasibinomial family taken to be 1.230381)<br>Null deviance: 73.505 on 62 degrees of freedom<br>Residual deviance: 70.059 on 59 degrees of freedom</td></tr>
<tr><td colspan="5">Analysis of Deviance Table (Type II tests)</td></tr>
<tr><td>Variable</td><td>Sum Sq</td><td>Df</td><td>F value</td><td>P value</td></tr>
<tr><td>NUTRIENT</td><td>0.553</td><td>1</td><td>0.4493</td><td>0.5053</td></tr>
<tr><td>PRESERVATIVES</td><td>3.016</td><td>1</td><td>2.4514</td><td>0.1228</td></tr>
<tr><td>TOTAL.EGGS</td><td>0.184</td><td>1</td><td>0.1496</td><td>0.7003</td></tr>
<tr><td>residuals</td><td>72.592</td><td>59</td><td></td><td></td></tr>
</table>

**Table S15.** Effect of nutrient level, preservative presence and clustering proportion on hatched egg to adult viability. Model coefficients of a quasibinomial GLM.

| **Model formula**<br>Y5 ~ NUTRIENT + PRESERVATIVES + PROPORTION.CLUSTERED, family = quasibinomial, data = EXP1A.DATA.M1F | | | | |
|---|---|---|---|---|
| **Model coefficients (quasibinomial GLM)** | | | | |
| **Variable** | **Estimate** | **Std. Error** | **t value** | **Pr(>\|t\|)** |
| (Intercept) | 2.8333 | 0.4104 | 6.903 | 4.62e-09 *** |
| NUTRIENT standard | -0.1705 | 0.3093 | -0.551 | 0.5837 |
| PRESERVATIVESpresent | 0.8046 | 0.4525 | 1.778 | 0.0807 . |
| PROPORTION.CLUSTERED | -1.0156 | 0.8658 | -1.173 | 0.2457 |
| (Dispersion parameter for quasibinomial family taken to be 1.093209)<br>Null deviance: 67.203 on 60 degrees of freedom<br>Residual deviance: 63.535 on 57 degrees of freedom | | | | |

| **Analysis of Deviance Table (Type II tests)** | | | | |
|---|---|---|---|---|
| **Variable** | **Sum Sq** | **Df** | **F value** | **P value** |
| NUTRIENT | 0.332 | 1 | 0.3039 | 0.58360 |
| PRESERVATIVES | 3.176 | 1 | 2.9052 | 0.09374 . |
| PROPORTION.CLUSTERED | 1.526 | 1 | 1.3961 | 0.24229 |
| residuals | 62.313 | 57 | | |

**Table S16.** Summary statistics for total eggs laid on substrates containing different combinations of antimicrobial preservatives.

| **treatment** | **N** | **mean egg count** | **sd** | **se** | **ci** |
|---|---|---|---|---|---|
| Std | 30 | 50.23 | 19.63 | 3.58 | 7.33 |
| S-P | 30 | 37.10 | 16.51 | 3.01 | 6.16 |
| EtOH | 30 | 39.50 | 17.57 | 3.21 | 6.56 |
| S-N | 30 | 42.47 | 18.87 | 3.44 | 7.05 |
| NP | 30 | 36.73 | 17.92 | 3.27 | 6.69 |

**Table S17.** Effect of preservatives on total eggs laid, under sterile conditions.

<table>
<tr><td colspan="5">Model formula<br>TOTAL.EGGS ~ TREATMENT, data = EXP2.DATA, init.theta = 5.209123224, link = log</td></tr>
<tr><td colspan="5">Model coefficients (negative binomial)</td></tr>
<tr><td>Variable</td><td>Estimate</td><td>Std. Error</td><td>Z value</td><td>Pr(>|z|)</td></tr>
<tr><td>(Intercept)</td><td>3.92</td><td>0.08</td><td>46.61</td><td>< 2e-16 ***</td></tr>
<tr><td>TREATMENTS-P</td><td>-0.30</td><td>0.12</td><td>-2.53</td><td>0.011 *</td></tr>
<tr><td>TREATMENTEtOH</td><td>-0.24</td><td>0.12</td><td>-2.01</td><td>0.044 *</td></tr>
<tr><td>TREATMENTS-N</td><td>-0.17</td><td>0.12</td><td>-1.41</td><td>0.159</td></tr>
<tr><td>TREATMENTNP</td><td>-0.31</td><td>0.12</td><td>-2.61</td><td>0.009 **</td></tr>
<tr><td colspan="5">(Dispersion parameter for Negative Binomial (5.21) family taken to be 1)<br>Null deviance: 166.44 on 149 degrees of freedom<br>Residual deviance: 156.98 on 145 degrees of freedom</td></tr>
</table>

<table>
<tr><td colspan="4">Analysis of Deviance Table (Type II tests)</td></tr>
<tr><td>Variable</td><td>Deviance</td><td>Df</td><td>P value</td></tr>
<tr><td>TREATMENT</td><td>9.46</td><td>4</td><td>0.051 .</td></tr>
</table>

**Table S18.** Summary statistics for the clustering proportion of eggs laid on substrates containing different combinations of antimicrobial preservatives.

| **treatment** | **N** | **mean clustering proportion** | **sd** | **se** | **ci** |
|---|---|---|---|---|---|
| Std | 30 | 0.09 | 0.07 | 0.01 | 0.03 |
| S-P | 30 | 0.08 | 0.08 | 0.01 | 0.03 |
| EtOH | 30 | 0.13 | 0.14 | 0.03 | 0.05 |
| S-N | 30 | 0.09 | 0.09 | 0.02 | 0.03 |
| NP | 30 | 0.07 | 0.07 | 0.01 | 0.03 |

**Table S19.** Effect of preservatives and total eggs on clustering proportion, under sterile conditions.

| **Model formula**<br>Y2B ~ TREATMENT + TOTAL.EGGS, family = quasibinomial, data = EXP2.DATA | | | | |
|---|---|---|---|---|
| **Model coefficients (quasibinomial GLM)** | | | | |
| **Variable** | **Estimate** | **Std. Error** | **t value** | **Pr(>\|t\|)** |
| (Intercept) | -3.20 | 0.30 | -10.56 | < 2e-16 *** |
| TREATMENTS-P | 0.10 | 0.24 | 0.44 | 0.660 |
| TREATMENTEtOH | 0.45 | 0.21 | 2.13 | 0.035 * |
| TREATMENTS-N | 0.12 | 0.21 | 0.57 | 0.569 |
| TREATMENTNP | -0.06 | 0.24 | -0.24 | 0.811 |
| TOTAL.EGGS | 0.02 | 0.00 | 4.01 | 9.81e-05 *** |
| (Dispersion parameter for quasibinomial family taken to be 2.84)<br>Null deviance: 471.46 on 149 degrees of freedom<br>Residual deviance: 408.17 on 144 degrees of freedom | | | | |
| **Analysis of Deviance Table (Type II tests)** | | | | |
| **Variable** | **Sum Sq** | **Df** | **F value** | **P value** |
| TREATMENT | 17.88 | 4 | 1.57 | 0.185 |
| TOTAL.EGGS | 46.68 | 1 | 16.41 | 8.31e-05 *** |
| residuals | 409.67 | 144 | | |

**Table S20.** Summary statistics for egg to adult viability of eggs laid on substrates containing different combinations of antimicrobial preservatives.

| **treatment** | **N** | **mean egg to adult viability** | **sd** | **se** | **ci** |
|---|---|---|---|---|---|
| Std | 27 | 0.88 | 0.14 | 0.03 | 0.06 |
| S-P | 29 | 0.90 | 0.10 | 0.02 | 0.04 |
| EtOH | 30 | 0.89 | 0.10 | 0.02 | 0.04 |
| S-N | 30 | 0.88 | 0.10 | 0.02 | 0.04 |
| NP | 30 | 0.88 | 0.10 | 0.02 | 0.04 |

**Table S21.** Effect of preservative treatment and clustering proportion on egg to adult viability.

<table>
<tr><td colspan="5">Model formula<br>Y2C ~ TREATMENT + PROPORTION.CLUSTERED, family = quasibinomial, data = EXP2.DATA.M2C</td></tr>
<tr><td colspan="5">Model coefficients (quasibinomial GLM)</td></tr>
<tr><td>Variable</td><td>Estimate</td><td>Std. Error</td><td>t value</td><td>Pr(>|t|)</td></tr>
<tr><td>(Intercept)</td><td>2.17</td><td>0.22</td><td>10.10</td><td><2e-16 ***</td></tr>
<tr><td>TREATMENTS-P</td><td>0.00</td><td>0.28</td><td>0.01</td><td>0.990</td></tr>
<tr><td>TREATMENTEtOH</td><td>0.07</td><td>0.27</td><td>0.27</td><td>0.787</td></tr>
<tr><td>TREATMENTS-N</td><td>-0.10</td><td>0.26</td><td>-0.40</td><td>0.691</td></tr>
<tr><td>TREATMENTNP</td><td>-0.07</td><td>0.27</td><td>-0.25</td><td>0.802</td></tr>
<tr><td>PROPORTION.CLUSTERED</td><td>-1.05</td><td>1.08</td><td>-0.98</td><td>0.329</td></tr>
<tr><td colspan="5">(Dispersion parameter for quasibinomial family taken to be 4.48)<br>Null deviance: 530.87 on 145 degrees of freedom<br>Residual deviance: 524.92 on 140 degrees of freedom</td></tr>
<tr><td colspan="5">Analysis of Deviance Table (Type II tests)</td></tr>
<tr><td>Variable</td><td>Sum Sq</td><td>Df</td><td>F value</td><td>P value</td></tr>
<tr><td>TREATMENT</td><td>2.31</td><td>4</td><td>0.13</td><td>0.972</td></tr>
<tr><td>PROPORTION.CLUSTERED</td><td>4.19</td><td>1</td><td>0.94</td><td>0.335</td></tr>
<tr><td>residuals</td><td>627.41</td><td>140</td><td></td><td></td></tr>
</table>

**Table S22.** Effect of preservative treatment and total eggs on egg to adult viability.

<table>
<tr><td colspan="5">Model formula<br>Y2C ~ TREATMENT + TOTAL.EGGS, family = quasibinomial, data = EXP2.DATA.M2C</td></tr>
<tr><td colspan="5">Model coefficients (quasibinomial GLM)</td></tr>
<tr><td>Variable</td><td>Estimate</td><td>Std. Error</td><td>t value</td><td>Pr(>|t|)</td></tr>
<tr><td>(Intercept)</td><td>1.95</td><td>0.33</td><td>5.94</td><td>2.15e-08 ***</td></tr>
<tr><td>TREATMENTS-P</td><td>0.04</td><td>0.28</td><td>0.14</td><td>0.893</td></tr>
<tr><td>TREATMENTEtOH</td><td>0.06</td><td>0.27</td><td>0.21</td><td>0.831</td></tr>
<tr><td>TREATMENTS-N</td><td>-0.10</td><td>0.26</td><td>-0.38</td><td>0.703</td></tr>
<tr><td>TREATMENTNP</td><td>-0.03</td><td>0.27</td><td>-0.10</td><td>0.917</td></tr>
<tr><td>TOTAL.EGGS</td><td>0.00</td><td>0.01</td><td>0.43</td><td>0.671</td></tr>
<tr><td colspan="5">(Dispersion parameter for quasibinomial family taken to be 4.39)<br>Null deviance: 530.87 on 145 degrees of freedom<br>Residual deviance: 528.32 on 140 degrees of freedom</td></tr>
<tr><td colspan="5">Analysis of Deviance Table (Type II tests)</td></tr>
<tr><td>Variable</td><td>Sum Sq</td><td>Df</td><td>F value</td><td>P value</td></tr>
<tr><td>TREATMENT</td><td>1.88</td><td>4</td><td>0.107</td><td>0.980</td></tr>
<tr><td>TOTAL.EGGS</td><td>0.80</td><td>1</td><td>0.182</td><td>0.670</td></tr>
<tr><td>residuals</td><td>614.68</td><td>140</td><td></td><td></td></tr>
</table>

**Table S23.** Summary statistics for total eggs laid on substrates containing different microbial communities.

| treatment | N | mean eggs | sd | se | ci |
|---|---|---|---|---|---|
| none | 30 | 57.03 | 30.13 | 5.50 | 11.25 |
| filtered | 30 | 54.23 | 31.34 | 5.72 | 11.70 |
| commensal | 30 | 56.00 | 31.88 | 5.82 | 11.90 |
| A. faecalis | 30 | 59.53 | 36.09 | 6.59 | 13.47 |

**Table S24.** Effect of microbial community on total eggs laid.

<table>
<tr><td colspan="5">Model formula<br>TOTAL.EGGS ~ TREATMENT, data = EXP3.DATA, init.theta = 2.117455871, link = log</td></tr>
<tr><td colspan="5">Model coefficients (negative binomial GLM)</td></tr>
<tr><td>Variable</td><td>Estimate</td><td>Std. Error</td><td>t value</td><td>Pr(>|t|)</td></tr>
<tr><td>(Intercept)</td><td>4.04</td><td>0.13</td><td>31.65</td><td><2e-16 ***</td></tr>
<tr><td>TREATMENTfiltered</td><td>-0.05</td><td>0.18</td><td>-0.28</td><td>0.781</td></tr>
<tr><td>TREATMENTcommensal</td><td>-0.02</td><td>0.18</td><td>-0.10</td><td>0.919</td></tr>
<tr><td>TREATMENTA.faecalis</td><td>0.04</td><td>0.18</td><td>0.24</td><td>0.812</td></tr>
<tr><td colspan="5">(Dispersion parameter for Negative Binomial (2.1175) family taken to be 1)<br>Null deviance: 143.14 on 119 degrees of freedom<br>Residual deviance: 142.86 on 116 degrees of freedom</td></tr>
<tr><td colspan="5">Analysis of Deviance Table (Type II tests)</td></tr>
<tr><td>Variable</td><td>LR Chisq</td><td>Df</td><td colspan="2">P value</td></tr>
<tr><td>TREATMENT</td><td>0.28</td><td>3</td><td colspan="2">0.964</td></tr>
</table>

**Table S25.** Summary statistics for clustering proportion of eggs laid on substrates containing different microbial communities.

| treatment | N | mean clustering proportion | sd | se | ci |
|---|---|---|---|---|---|
| none | 30 | 0.14 | 0.11 | 0.02 | 0.04 |
| filtered | 28 | 0.14 | 0.13 | 0.02 | 0.05 |
| commensal | 28 | 0.13 | 0.09 | 0.02 | 0.03 |
| A. faecalis | 28 | 0.16 | 0.13 | 0.02 | 0.05 |

**Table S26.** Effect of microbial community and total eggs on egg clustering proportion.

| **Model formula**<br>Y3B ~ TREATMENT + TOTAL.EGGS, family = quasibinomial, data = EXP3.DATA | | | | |
|---|---|---|---|---|
| **Model coefficients (quasibinomial GLM)** | | | | |
| **Variable** | **Estimate** | **Std. Error** | **t value** | **Pr(>\|t\|)** |
| (Intercept) | -2.14 | 0.23 | -9.04 | 6.18e-15 *** |
| TREATMENTfiltered | -0.05 | 0.22 | -0.25 | 0.806 |
| TREATMENTcommensal | -0.17 | 0.22 | -0.79 | 0.431 |
| TREATMENTA.faecalis | 0.10 | 0.21 | 0.48 | 0.636 |
| TOTAL.EGGS | 0.01 | 0.00 | 2.82 | 0.006 ** |
| (Dispersion parameter for quasibinomial family taken to be 5.18)<br>Null deviance: 631.98 on 114 degrees of freedom<br>Residual deviance: 578.03 on 110 degrees of freedom | | | | |
| **Analysis of Deviance Table (Type II tests)** | | | | |
| **Variable** | **Sum Sq** | **Df** | **F value** | **P value** |
| TREATMENT | 8.73 | 3 | 0.562 | 0.641 |
| TOTAL.EGGS | 41.12 | 1 | 7.945 | 0.006 ** |
| residuals | 569.35 | 110 | | |

**Table S27.** Summary statistics for egg to adult viability of eggs laid on substrates containing different microbial communities.

| **treatment** | **N** | **mean egg to adult viability** | **sd** | **se** | **ci** |
|---|---|---|---|---|---|
| none | 30 | 0.88 | 0.10 | 0.02 | 0.04 |
| filtered | 28 | 0.88 | 0.12 | 0.02 | 0.05 |
| commensal | 28 | 0.90 | 0.08 | 0.01 | 0.03 |
| A. faecalis | 29 | 0.87 | 0.18 | 0.03 | 0.07 |

**Table S28.** Effect of microbial community and egg clustering proportion on egg to adult viability.

**Model formula**

Y3C ~ TREATMENT + PROPORTION.CLUSTERED, family = quasibinomial, data = EXP3.DATA

**Model coefficients (quasibinomial GLM)**

| Variable | Estimate | Std. Error | t value | Pr(>\|t\|) |
|---|---|---|---|---|
| (Intercept) | 2.15 | 0.20 | 10.64 | <2e-16 *** |
| TREATMENTfiltered | -0.08 | 0.23 | -0.36 | 0.723 |
| TREATMENTcommensal | -0.02 | 0.23 | -0.08 | 0.934 |
| TREATMENTA.faecalis | 0.01 | 0.23 | 0.04 | 0.971 |
| PROPORTION.CLUSTERED | 0.07 | 0.75 | 0.09 | 0.930 |

(Dispersion parameter for quasibinomial family taken to be 4.14)
Null deviance: 415.41 on 113 degrees of freedom
Residual deviance: 414.57 on 109 degrees of freedom

**Analysis of Deviance Table (Type II tests)**

| Variable | Sum Sq | Df | F value | P value |
|---|---|---|---|---|
| TREATMENT | 0.77 | 3 | 0.06 | 0.980 |
| PROPORTION.CLUSTERED | 0.03 | 1 | 0.01 | 0.930 |
| residuals | 450.49 | 109 | | |

**Table S29.** Effect of microbial community and total eggs on egg to adult viability.

**Model formula**

Y3C ~ TREATMENT + TOTAL.EGGS, family = quasibinomial, data = EXP3.DATA

**Model coefficients (quasibinomial GLM)**

| Variable | Estimate | Std. Error | t value | Pr(>\|t\|) |
|---|---|---|---|---|
| (Intercept) | 2.11 | 0.25 | 8.54 | 8.54e-14 *** |
| TREATMENTfiltered | -0.08 | 0.23 | -0.36 | 0.724 |
| TREATMENTcommensal | -0.02 | 0.23 | -0.09 | 0.926 |
| TREATMENTA.faecalis | -0.00 | 0.23 | -0.01 | 0.991 |
| TOTAL.EGGS | 0.00 | 0.00 | 0.32 | 0.748 |

(Dispersion parameter for quasibinomial family taken to be 4.14)
Null deviance: 419.91 on 114 degrees of freedom
Residual deviance: 418.73 on 110 degrees of freedom

**Analysis of Deviance Table (Type II tests)**

| Variable | Sum Sq | Df | F value | P value |
|---|---|---|---|---|
| TREATMENT | 0.68 | 3 | 0.05 | 0.983 |
| TOTAL.EGGS | 0.43 | 1 | 0.10 | 0.748 |
| residuals | 456.19 | 110 | | |

### Supplementary Figures

| | | | |
|---|---|---|---|
| | 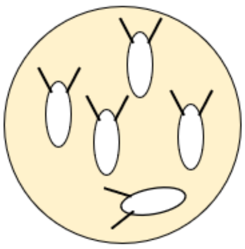 | 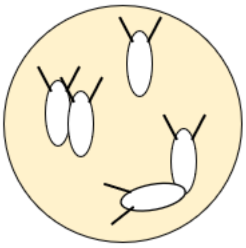 | 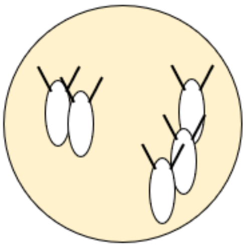 |
| Number of eggs in clusters (C) | 0 | 4 | 5 |
| Total eggs (N) | 5 | 5 | 5 |
| Clustering proportion (C / N) | 0 | 0.8 | 1 |

**Figure S1. Example egg clusters and calculation of egg clustering proportion.** From left to right, the schematics in the top row show an example where zero eggs are clustered, four eggs are clustered (2 clusters of 2 eggs) and 5 eggs are clustered (2 clusters of 2 and 3 eggs).

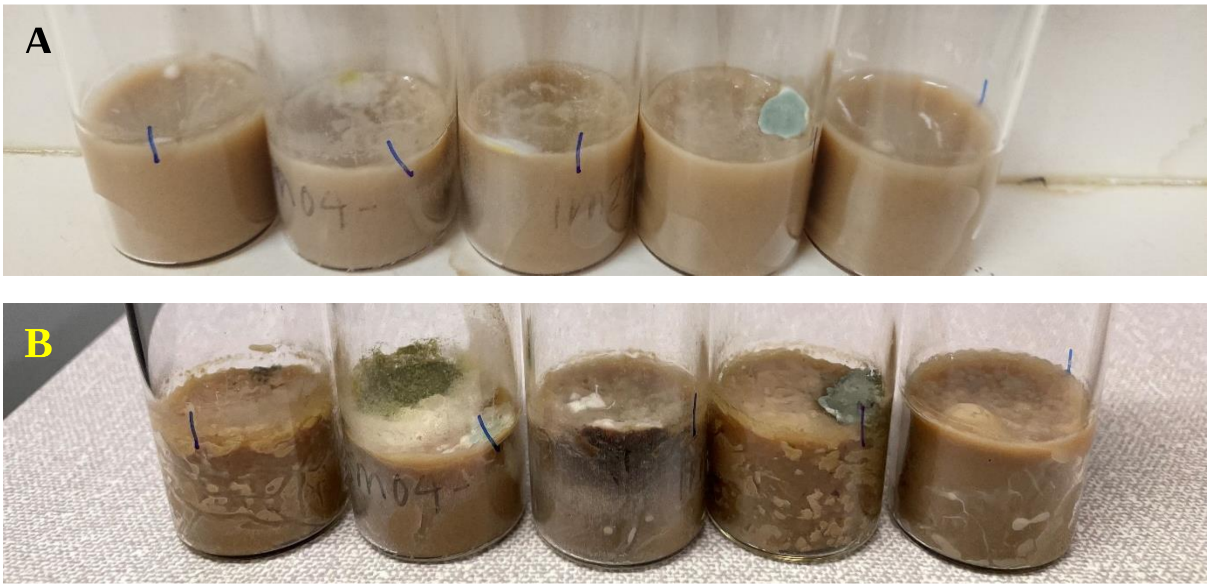


**Figure S2. Microbial growth on oviposition substrates lacking antimicrobial preservatives.** Photos were taken 48h (A) or 96h (B) following the oviposition assay. The same vials are photographed in the same order in both photos.

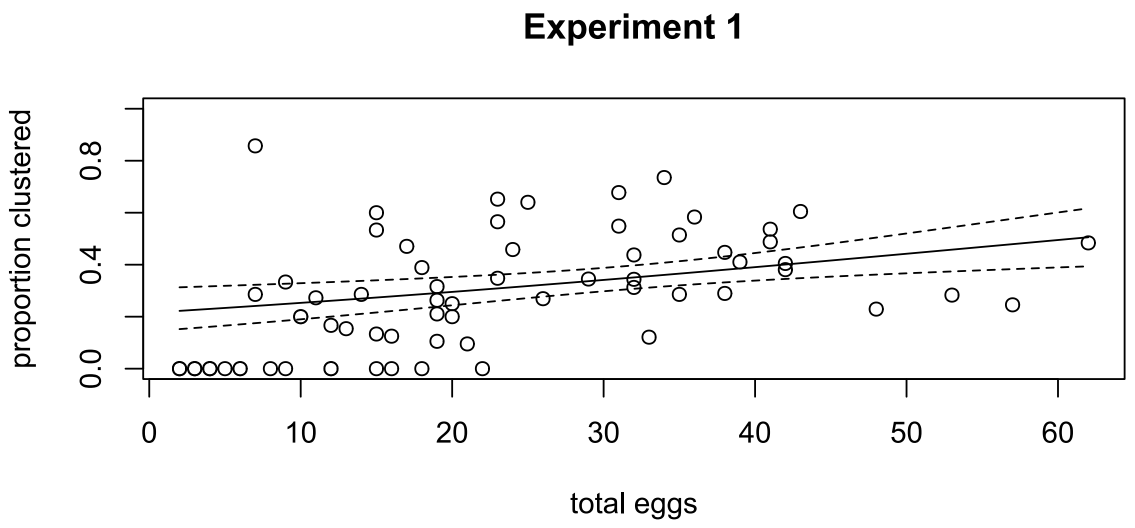


**Figure S3.** Relationship between the total number of eggs in a vial and the proportion of those eggs that are in clusters from Experiment 1 (Effect of environmental microbes and nutrients on oviposition). The solid line represents the results of a generalised linear model with total eggs as the sole explanatory variable. Dashed lines indicate 95% confidence intervals.

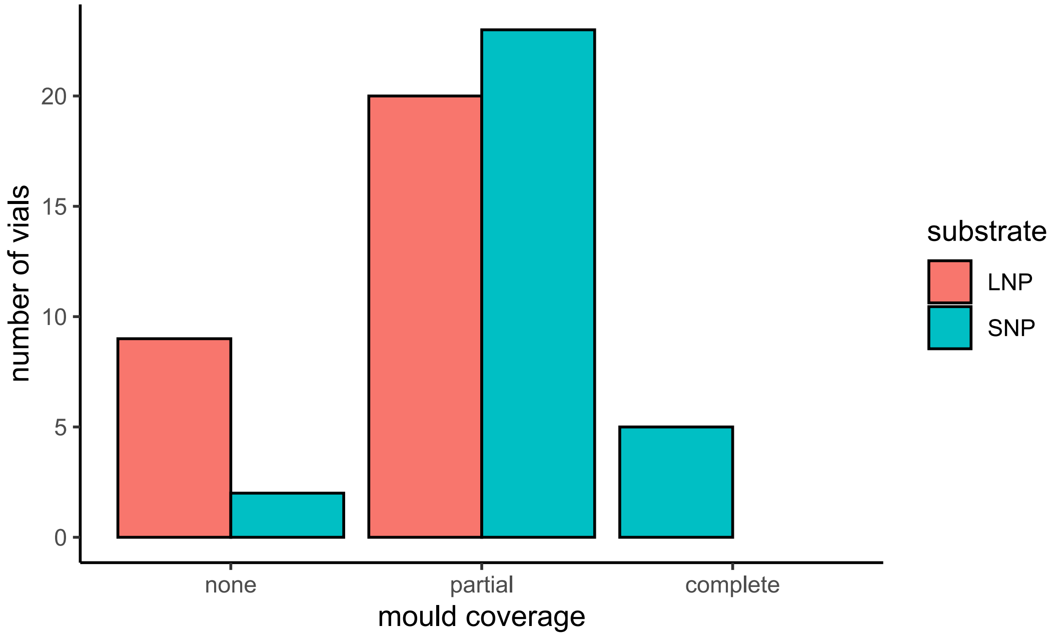


**Figure S4. Extent of microbial growth on the surface of substrates lacking preservatives, 24h following oviposition.** Substrates were either low nutrient (LNP) or standard nutrient (SNP). Microbial growth coverage on the surface of each substrate was visually inspected under a microscope and each substrate was scored as belonging to one of three categories – none (no microbial growth visible), partial (microbial growth visible, but across entire substrate) or complete (microbial growth completely obscuring underlying substrate).

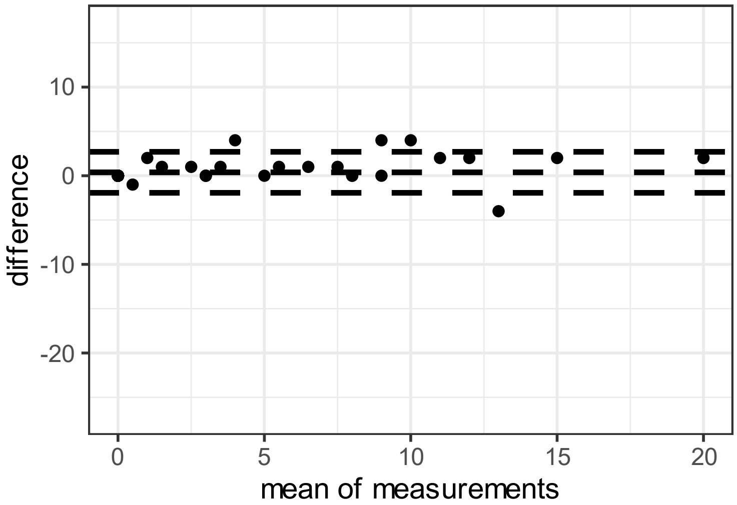


**Figure S5. Bland-Altman mean-difference plot showing consistency in identifying focal eggs in each vial, regardless of egg density.** The y axis is the difference between the number of focal eggs scored and the number of focal offspring scored. A difference of 0 indicates perfect agreement between eggs and offspring, negative differences indicate more offspring than eggs were scored, and a positive difference indicates fewer offspring than eggs. The x axis shows the mean of the two measurements (eggs and offspring). Each data point represents one vial. Dotted horizontal lines represent the mean difference and 95% confidence intervals.

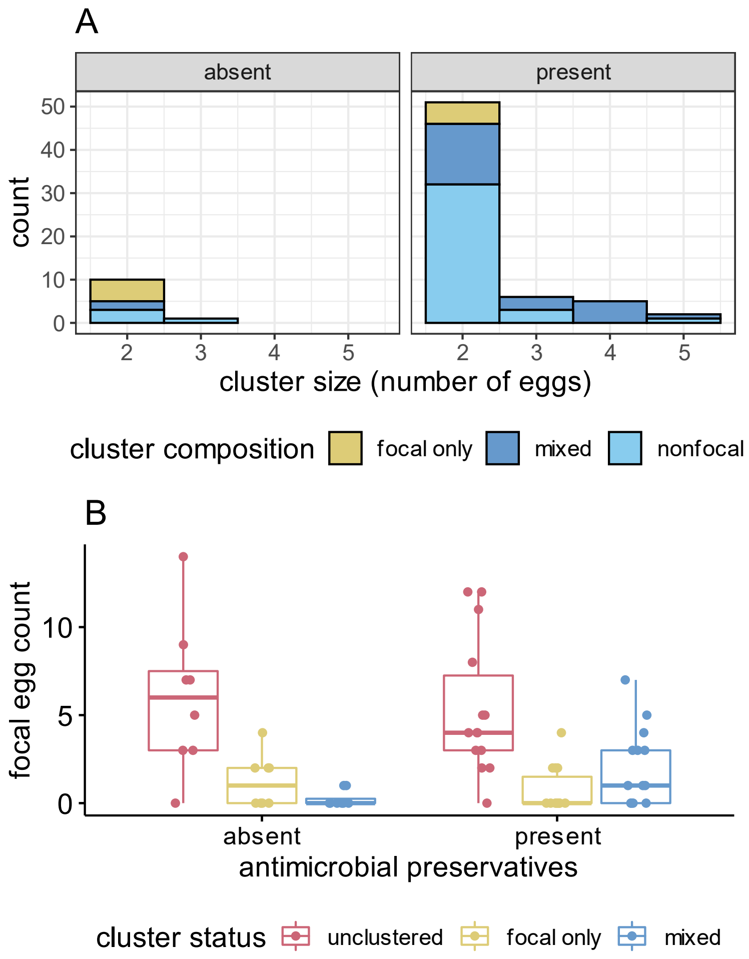


**Figure S6. Females lay eggs in mixed maternity clusters.** Clusters of eggs were counted from across oviposition substrates that either lacked preservatives ("absent") or contained preservatives ("present"), n = 30. One focal female and three non-focal females were laying on each substrate. Eggs from non-focal females were indistinguishable from one another. (A) shows the total number of clusters of different sizes from across all substrates, split by whether the cluster contained focal eggs only (yellow), had a mix of focal and non-focal eggs (dark blue) or had eggs laid by non-focal females only (light blue). Bars are stacked and non-overlapping. (B) shows the number of focal eggs on each substrate that were categorised as "unclustered" (single eggs not physically in contact with any other egg), "focal only" (a cluster containing only focal eggs) or "mixed" (a cluster of eggs containing ≥1 focal egg and ≥1 non-focal egg). Boxplots show the interquartile range (IQR) and median in the box, and whiskers represent the largest and smallest values within 1.5 times the IQR above and below the $75^{th}$ and $25^{th}$ percentiles, respectively. Raw data points are plotted with jitter.

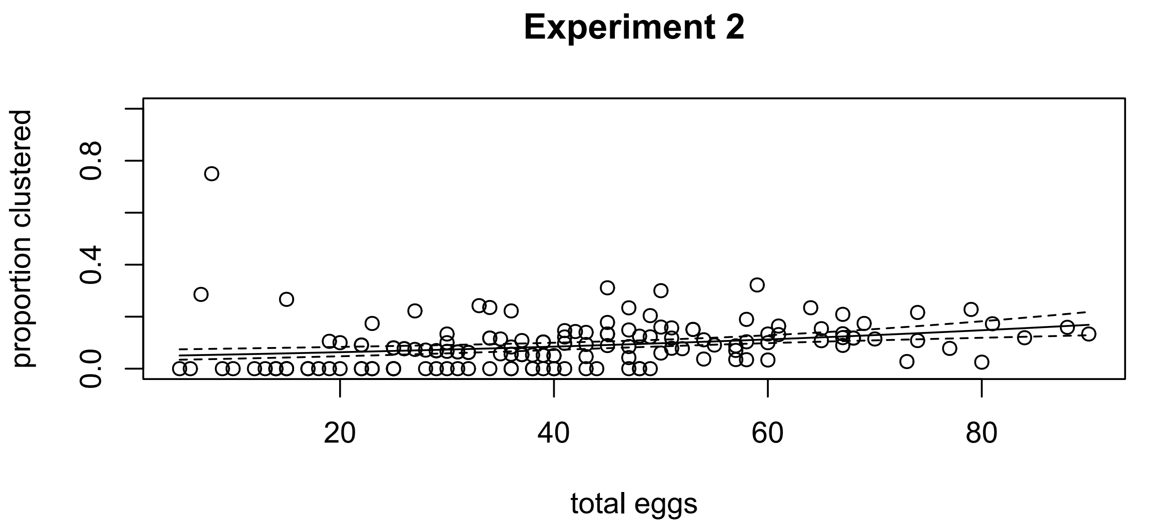


**Figure S7.** Relationship between the total number of eggs in a vial and the proportion of those eggs that are in clusters from Experiment 2 (Effect of antimicrobial preservatives on oviposition). The solid line represents the results of a generalised linear model with total eggs as the sole explanatory variable. Dashed lines indicate 95% confidence intervals.

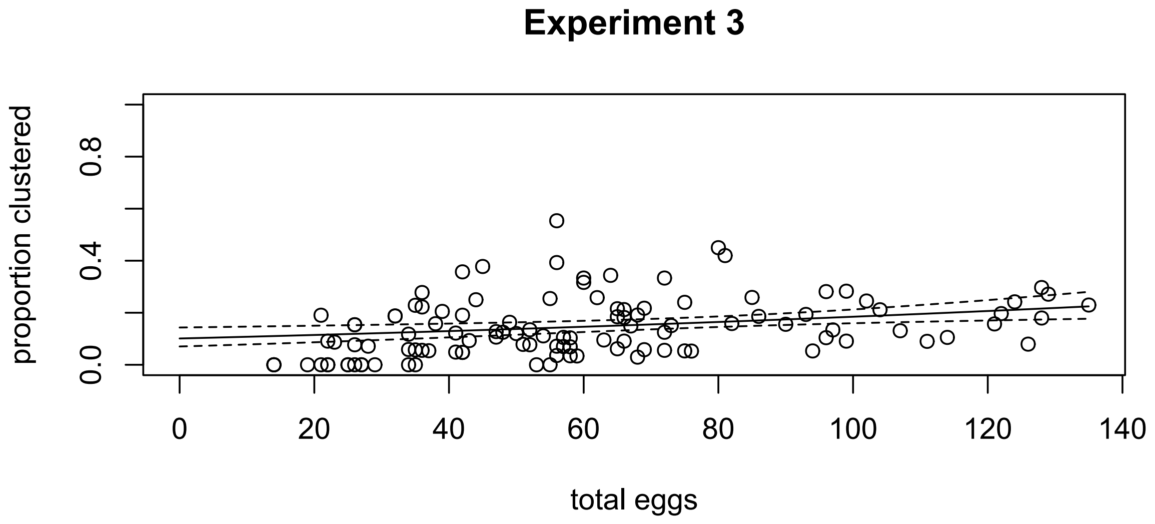


**Figure S8.** Relationship between the total number of eggs in a vial and the proportion of those eggs that are in clusters from Experiment 3 (Effect of commensal and pathogenic microbes on oviposition). The solid line represents the results of a generalised linear model with total eggs as the sole explanatory variable. Dashed lines indicate 95% confidence intervals.

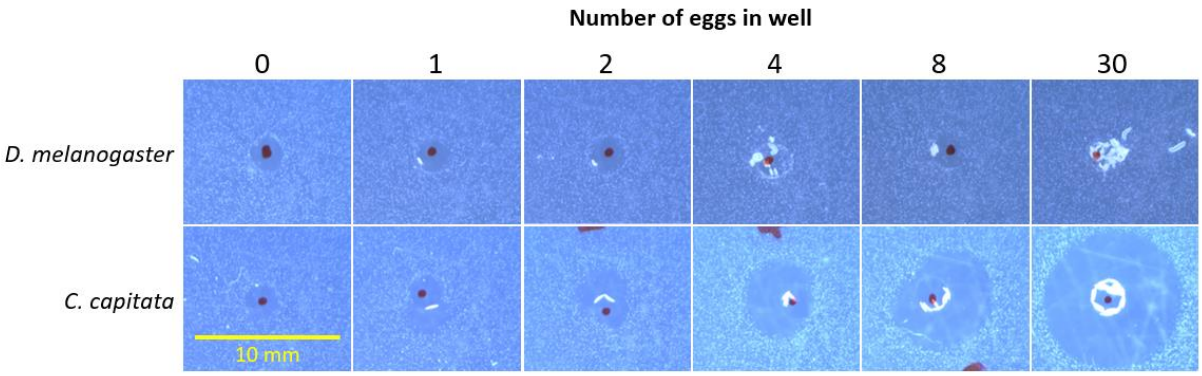


**Figure S9. Whole eggs of *D. melanogaster* do not exhibit antimicrobial activity against *E. coli*.** Intact, freshly laid eggs of either *D. melanogaster* (top row) or *C. capitata* (bottom row) were transferred to wells of LB agarose plates containing *E. coli* culture. Numbers of eggs used ranged from 0 to 30 (indicated above each column). Wells showing any surrounding zones of *E. coli* growth inhibition were photographed and arranged into the composite image shown. A few larvae that had hatched from the *D. melanoga*ster eggs during the overnight incubation are visible to the right of the well when 30 eggs were used. A scale bar of 10 mm is shown on the bottom left.
